# Drought to deluge: Differential impacts of snow on mountain chickadee reproduction across the Sierra Nevada mountains

**DOI:** 10.64898/2026.05.02.722414

**Authors:** Joseph F. Welklin, Lauren E. Whitenack, Benjamin R. Sonnenberg, Carrie L. Branch, Angela M. Pitera, Sofia M. Haley, Ai Ana Richmond, Vladimir V. Pravosudov

## Abstract

Changing climates are reshaping animal populations, but our understanding of how demographic trends are shaped by individual responses to local environmental conditions has historically been limited to long-term studies with restricted spatial scales. Participatory science (i.e. citizen science) programs have improved our ability to identify geographic variation in population-level trends, but these programs often lack insight into the individual-level processes that underly such variation. Here we present a new research framework that uses long-term data to validate participatory science observations as measurements of reproductive success, then explores reproductive responses to environmental variation across large geographic scales. We use this approach to investigate how reproduction in a montane songbird, the mountain chickadee (*Poecile gambeli*), varies in response to extreme scarcity and extreme accumulation of snow throughout the Sierra Nevada Mountains in North America. Reproductive responses to snowfall extremes varied considerably across elevations and latitudes. In the northern Sierra Nevada, mountain chickadee reproductive performance was negatively affected by snow deluge conditions at high elevations, whereas snow drought conditions reduced reproductive output at low elevations. In the central Sierras, drought conditions negatively impacted both elevations, but deluge conditions improved reproductive performance at both low and high elevations. Reproduction in the southern Sierra Nevada was nearly unaffected by spring snow levels. These variable responses to extreme snow accumulation across environments corroborate increasing evidence that biological responses to environmental variation are often nonstationary across spatial scales. Our results show that long-term studies can be used to inform and interpret participatory science data to provide a framework that strengthens our understanding of how individual behaviors underly differential responses to environmental variation across geographic scales.

## Introduction

Climatic extremes can have outsized impacts on animal reproduction (Ropert-Coudert et al., 2015; Wingfield et al., 2017), at times influencing long-term population patterns (Brawn et al., 2017; Grant et al., 2017). Extreme climate events are increasingly common (Cohen et al., 2018; Ummenhofer & Meehl, 2017) and while the effects of extreme temperature and rainfall on demographic processes are well-studied (Boyle et al., 2020; Dougherty et al., 2024), the effects of snowfall extremes on reproduction are less understood (Slatyer et al., 2022; Williams et al., 2015). Like rain, snowfall extremes can be characterized by an over-abundance or an exceptional lack of precipitation and may be direct or indirect. Extreme accumulation of snowfall can directly affect animal reproduction when out-of-season snowstorms prevent reproduction (Astheimer et al., 1995; Descamps et al., 2023), or when long-lasting snow cover delays breeding (Bison et al., 2020; Hendricks 2003; Kozlovsky et al., 2018; Whitenack et al., 2023) or hinders access to food resources (Schindler et al., 2026; Schmidt et al., 2019; White et al., 2025). Heavy snowpack may also indirectly affect reproduction by shifting the phenology of food resources, such as delaying the emergence of insect prey (Finn & Poff, 2008; Leingaertner et al., 2014). In contrast, an extreme lack of snowfall can indirectly affect reproduction in climates where winter snowfall is a major source of annual water (Van Horne et al., 1997). For example, in semi-arid habitats, a reduction in winter snow accumulation is associated with advanced phenology and decreased survival of the plants and insects that make up the basis of the vertebrate food web (Brunet et al., 2025; Forister et al., 2018; Halsch et al., 2024).

Understanding how these opposite snowfall extremes contribute to variation in animal reproduction across years and environments is critical for predicting future responses to environmental change and implementing conservation action.

The majority of the mountain west of the United States and its complex landscapes depend on accumulated winter snowfall (i.e., snowpack) for precipitation (Hale et al., 2023; Leung & Qian, 2009; Li et al., 2017), yet winters in the western United States are far from predictable. In some years the Sierra Nevada mountains experience very little snowfall (i.e. drought years), whereas other years are defined by extremely high snow accumulation (i.e. deluge years; Marshall et al., 2024), often a result of atmospheric rivers that deliver tremendous amounts of moisture from the tropics to the western United States (Dong et al., 2019; Margulis et al., 2016a). Recently, both drought and deluge are increasingly frequent in the Sierra Nevada (Diffenbaugh et al., 2015; Huning & AghaKouchak, 2020; Swain et al., 2018), but the magnitude of these extremes varies across elevations and latitudes. For example, the same storm system can cause flooding at lower elevations where temperatures are usually warmer yet bring over a meter of snow to high elevations where temperatures are typically colder (Dettinger, 2013; Leung & Qian, 2009). More broadly, southern latitudes typically receive less snowfall than northern latitudes (Huning & AghaKouchak, 2018). As a result, drought conditions at lower elevations and lower latitudes often correspond to average snowpack conditions at high elevations and higher latitudes in the same year, leading to differential selection on animal reproduction across populations inhabiting different elevations and latitudes (e.g. Forister et al., 2018).

A limitation of many current studies is the inability to measure these differential effects of climate extremes on reproduction across environments. Accurately measuring vertebrate reproduction is a time-intensive process, often only accomplished by long-term studies that focus on single populations. Long-term studies have generated irreplaceable insights into how individual animals respond to yearly environmental variation and how these responses affect population dynamics (e.g. Radchuk et al., 2026), but the range of environmental extremes experienced by single populations is often restricted compared to the environmental variation a species experiences across its range. Thus, our knowledge of how species respond to climatic extremes across different environments is frequently limited to extrapolation and inference.

Participatory science programs (sometimes called citizen or community science), structured point counts, and other targeted survey efforts are closing these knowledge gaps (Evans et al., 2024; Fink et al., 2020; Senzaki et al., 2020; Socolar et al., 2025; Youngflesh et al., 2023), but the broad-scale nature of these data sources typically limits researchers’ abilities to identify the individual-level responses that contribute to population-level trends. Here we propose a new framework for investigating animal responses to environmental variation that first validates participatory science data using long-term data, then explores the corroborated participatory science data on a much larger geographic scale to investigate differences in population-level responses to environmental extremes.

Our focal species is the mountain chickadee (*Poecile gambeli*), a small, non-migratory songbird that breeds throughout the mountain west of the United States, including the Sierra Nevada range. Mountain chickadees are well-adapted to year-round montane living (Pravosudov et al., 2025; Sonnenberg et al., 2025), but recent drought and deluge snowfall extremes have strongly impacted their reproductive patterns. Our long-term study revealed that the 2012-2015 drought – among the most severe in the in the Sierra Nevada in the last 1000 years (Griffin & Anchukaitis, 2014) – resulted in fewer chickadee offspring fledged at lower elevations compared to summers following average snow conditions. Yet, reproduction at high elevations was not affected by the drought. In contrast, the snow deluge conditions that followed in spring 2017 and spring 2019 negatively affected reproductive output at higher elevations, but not lower elevations (Kozlovsky et al., 2018; Whitenack et al., 2023). This disparity in reproduction occurred despite our low and high elevation study sites being separated by only 3.5km in distance and ca. 500m in altitude. These sites occur across a snow line, above which winter precipitation is more likely to fall as snow which often starts earlier in autumn and persists well into the summer. Below the snow line, snow accumulation starts later in autumn and melts earlier in the spring (Kirchner et al., 2014; Minder et al., 2013). As a result, low snow years still result in substantial snow depth and average reproductive output at the high elevation site, but high snow years negatively affect reproduction at high elevations while boosting reproduction at low elevations when snow cover is sustained into the spring (Kozlovsky et al., 2018; Whitenack et al., 2023). The mechanisms underlying these differential reproductive responses to spring snow depth are still under investigation, but current findings indicate that drought conditions at low elevations reduce invertebrate abundance (Forister et al., 2018; Halsch et al., 2024), likely resulting in fewer chickadee offspring surviving to fledging due to reduced food availability. At high elevations, deluge conditions delay breeding, resulting in fewer eggs being laid and thus fewer offspring fledging from nests (Whitenack et al., 2023).

Like many long-term studies, our focal population represents an extremely small portion of the species’ overall range, meaning we are unsure how consistent these relationships between snowfall and reproduction are throughout the mountain west. In this study, we focus on the Sierra Nevada mountains, a region known for its extreme elevational gradients that occur across an extensive latitudinal range. We explore the effect of snow depth on mountain chickadee reproduction across the Sierras by using our long-term data to inform analyses of participatory science data. We first match yearly variation in offspring production in our 11-year database of breeding records to local post-breeding observations of chickadees in the eBird dataset (Sullivan et al., 2009). After finding support that yearly variation in breeding output is reflected in eBird observations, we explore how relationships between snowfall and mountain chickadee reproduction vary across the Sierra Nevada. Based on our long-term study, we predicted that chickadee reproduction at low elevations would be negatively affected by drought conditions, whereas reproduction at high elevations would be negatively affected by snow deluge conditions, but we were unsure how these relationships might change across latitudes that experience different snowfall and snowmelt regimes. This study synchronizes traditionally separate datasets to provide new insights into how variable environments and climatic extremes interact to differentially influence animal reproduction across large geographic scales.

## Methods

### Long-term study site

We monitored mountain chickadee reproduction in approximately 350 wooden nest boxes at two elevations from April through August across 11 years (2014-2024) at Sagehen Experimental Forest (Sagehen Creek Field Station, University of California, Berkeley), California, USA. Breeding typically begins in late April or early May, and most nests fledge before August. The low elevation site ranged from 1965m to 2070m (6447ft to 6791ft) elevation and included approximately 200 nest boxes, whereas the high elevation site ranged from 2380m to 2590m (7808ft to 8497ft) elevation and included approximately 150 nest boxes (the number of boxes varied slightly across years). We determined the brood size of each nest (the number of offspring in each nest) when nestlings were 16 days old (nestlings were considered 1 day old on hatch day). Fledgling typically occurs 20 to 24 days after hatching (Grundel, 1987), but our observations indicate that nearly all nestlings that reach 16 days of age successfully fledge their nests (VVP personal observation), thus brood size on day 16 is an accurate measure of how many offspring were produced in each nest (Welklin et al., 2025). Most nest boxes were protected by metal predator guard sheaths which reduced predation of eggs and nestlings by various species of chipmunks. As a result, nests were rarely predated (less than 2% of nests were renests after a previous failed nesting attempt) and failed nests were not included in observations of brood size, meaning our comparisons of brood sizes across years represent variation among successful nests, or relative reproduction across years. Secondary nests initiated after successfully fledging a first nest were rare and typically unsuccessful (Whitenack et al., 2023), so were excluded from analyses of brood size. All methods and procedures were approved by the University of Nevada Reno (UNR) Institutional Animal Care and Use Committee (IACUC) in accordance with UNR IACUC protocols 20-11-1103, 20-06-1014, 20-08-1062, under California Department of Fish and Wildlife Permit SC-193630001-20007-001.

### Snow depth and reproduction in the long-term study

We obtained yearly measurements of April 1 snow depth from SNOTEL (Snow Telemetry) sites maintained by the United States Department of Agriculture Natural Resources Conservation Service (https://www.nrcs.usda.gov) located within and near the long-term study site (see Supplemental Methods). Snow depth on April 1 is commonly used to compare yearly variation in snow depth in the Sierra Nevada mountains because prior to April 1, most precipitation falls as snow and rapid melting usually does not usually begin until later in the spring (Aguado, 1990; Hatchett & McEvoy, 2018; Margulis et al., 2016b; Marshall et al., 2024). In analyses of our long-term data, we consider years 2014 and 2015 to be ‘drought’ years based on the low April 1 snow depths at the low elevation site (Figure 1A), and we consider years 2017, 2019, and 2023 to be ‘deluge’ years based on the high April 1 snow depths at the high elevation site (Figure 1A). Drought conditions at the low elevation site did not necessarily reflect low-snow conditions at the high-elevation site and vice versa for deluge conditions at the high elevation site (Kozlovsky et al., 2018; Whitenack et al., 2023). Years 2014 and 2015 were the last 2 years in a 4-year millennium-level drought (Griffin & Anchukaitis, 2014), and years 2017, 2019, and 2023 are widely recognized as some of the largest snow years in the recent history of the Sierra Nevada (Marshall et al., 2024, Swain et al., 2018). All other years were considered average snow years.

**Figure 1.**
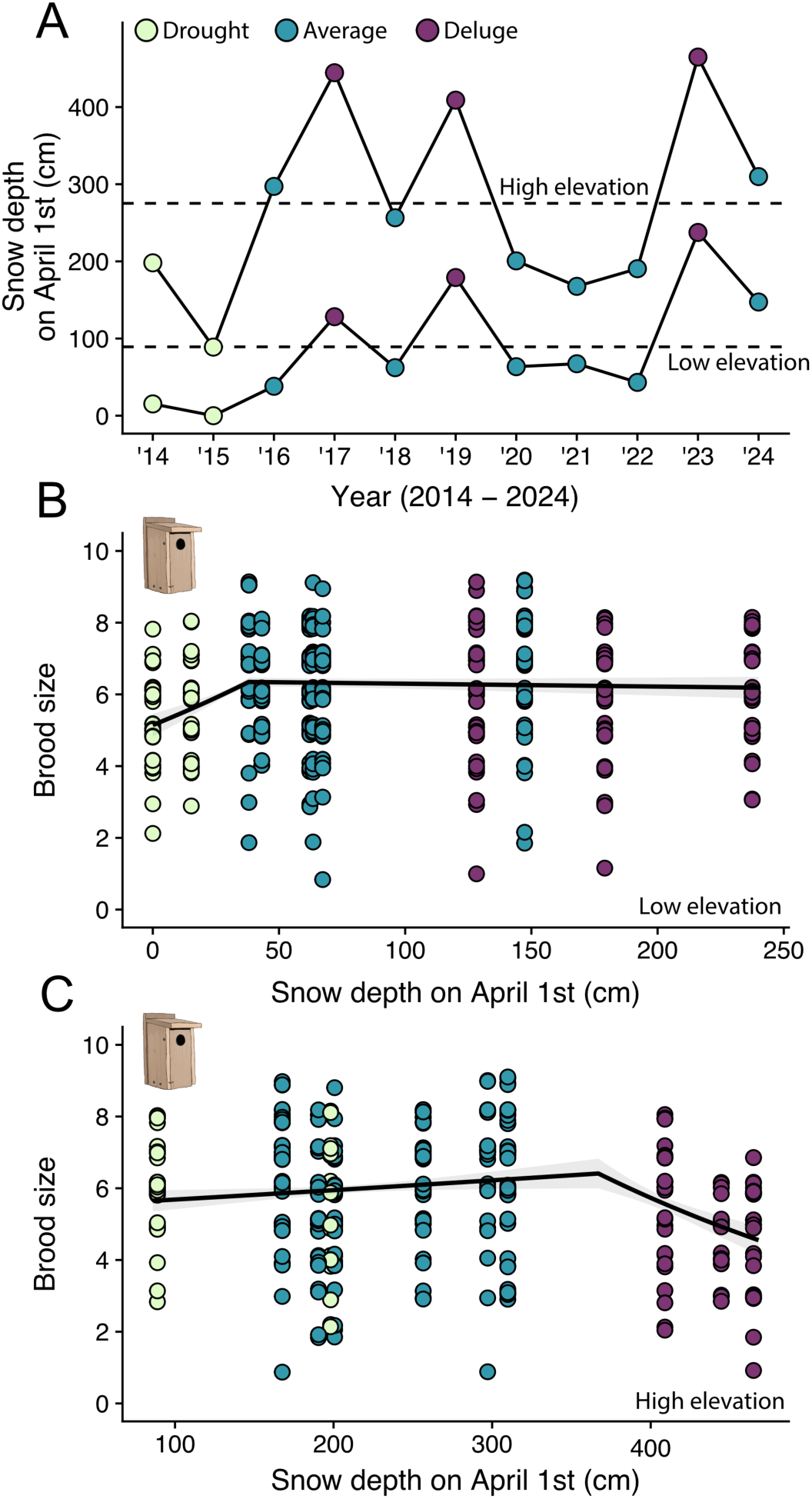
Relationships between snow depth and reproduction vary across elevations for mountain chickadees in the long-term study. A) Yearly snow depth on April 1^st^ at the high elevation site (2541m elevation; upper line) and yearly snow depth on April 1^st^ at the low elevation site (averaged from two sites: 1962m, 2128m elevation; lower line). Dotted lines show average snow depth from years 2014 through 2024 at each site. Points in all plots are colored relative to yearly snow depth categories. Drought years were based on April snow levels at the low-elevation site and deluge years were based on April snow levels at the high-elevation site. B) Relationship between brood size (number of offspring) and snow depth at the low elevation site (1965m – 2070m) in the long-term study. C) Relationship between brood size and snow depth at the high elevation site (2380m – 2590m) in the long-term study. Points in B and C are jittered slightly on the y-axis.

We modeled the relationship between snow depth and reproduction in the long-term study separately for the low and high-elevation sites because previous work has indicated that the variation in snow depths across elevations differentially affects reproduction (Kozlovsky et al., 2018; Whitenack et al., 2023). The effect of April 1 snow depth on brood size was modeled using generalized linear models with the Generalized Poisson distribution (Joe & Zhu, 2005) and a log link in the R package ‘glmmTMB’ (Brooks et al., 2017). We first modeled a categorical effect of snow conditions on brood size. We compared drought years to average and deluge years at the low elevation site and compared deluge years to average and drought years at the high elevation site. Next, we tested for a continuous effect of snow depth on chickadee reproduction. Likelihood ratio tests were used to compare whether models with linear, quadratic, or threshold effects of snow depth on April 1 best described variation in brood size. Threshold models are versions of linear or generalized linear models that allow the slope of the relationship between the response and a predictor to vary on either side of a specified breakpoint value (threshold) within the range of the predictor (e.g. Welklin et al., 2025). Here we explored whether the slope of the relationship between brood size and snow depth changed with increasing snow depth. We determined the best-supported snow depth threshold for each elevation by comparing AIC scores of models with breakpoints at 1cm increments of snow depth ranging from 5cm greater than the lowest snow depth value to 5cm less than the highest snow depth value. These constrained ranges helped to limit the effect of single extreme years and allowed us to compare support for general patterns that were supported by data from multiple years. For threshold models, we present the slopes and p-values for the lines before and after the threshold. All analyses were conducted in R version 4.4.2 (R Core Team 2024).

### eBird dataset

We downloaded observations of mountain chickadees (basic dataset) and sampling attempts (sampling dataset) from eBird.org for California and Nevada from July through September from 2014 through 2024 (eBird basic dataset 2024; Sullivan et al., 2009). We zero-filled complete checklists in which mountain chickadees were not reported and selected unique checklists using the ‘auk’ package in R (Strimas-Mackey et al., 2025). Observations were filtered based on a geographic perimeter that encompassed the main Sierra Nevada mountain range in California and Nevada (coordinates: southeast corner: 35.714° N, -116.426° W; southwest corner: 34.794° N, -118.450° W; northwest corner: 40.605° N, -122.477° W; northeast corner: 40.966° N, -120.719° W) and excluded the eastern Nevada mountain ranges that are less forested and typically less populated by mountain chickadees. We selected checklists with sampling periods that were at least 5 minutes long, but were 6 hours or less in length, were 7km or less in distance, had 10 or fewer observers, and were collected within local late summer daylight hours (after 6am PST and before 7:30pm PST). We were specifically interested in comparing mountain chickadee counts in areas where this species reliably occurs, so we assigned each checklist to a 5km x 5km grid cell and excluded cells in which more than 70% of the checklists in the cell did not report mountain chickadees. This step removed checklists from arid ecosystems like Mono Lake, a popular birding hotspot where mountain chickadees have been observed but are not regularly seen.

To compare eBird observations to the low elevation site (1965 - 2070m; 6447 - 6791ft), we filtered for checklists collected between 1829m and 2134m (6000-7000ft) elevation, and to compare eBird observations to the high elevation site (2380 - 2590m; 7808 - 8497ft), we filtered for checklists collected at or above 2377m (7800ft) elevation. More specific geographic regions were identified in later steps. Our goal was to use eBird data to measure reproductive output, so we filtered for eBird checklists collected after most initial nests had fledged in the long-term study: 99% of initial nests fledged by July 17^th^ (day 198 on a 365-day calendar) at the low elevation site and 99% of initial nests fledged by July 28^th^ (day 209 on a 365-day calendar) at the high elevation site (Figure S1). Since renests are rare (Whitenack et al., 2023), few nests fledge after these dates, so observations collected after these dates represent the post-fledging period when recently fledged juveniles were still dependent on their parents, were beginning to disperse, or had already dispersed and settled on a permanent territory (McCallum et al., 2020). Mountain chickadees disperse short distances (median distance approximately 700m; Whitenack et al., 2025), so post-dispersal eBird observations should be representative of local breeding production (see below for validation of this assumption). Chickadees are extremely vocal during this period and therefore should be easy for birders to detect. End dates were identified in later steps.

### Environmental data

We obtained information on environmental conditions in the Sierra Nevada mountains from multiple sources to help predict mountain chickadee abundance in eBird data. Elevation data was obtained for each checklist location using Shuttle Radar Topography Mission (SRTM; NASA JPL) Mapzen terrain tiles (Tilezen, 2025), courtesy of the U.S. Geological Survey and accessed using the ‘elevatr’ package in R (Hollister, 2025). We matched annual land cover data to each eBird checklist location using the Moderate Resolution Imaging Spectroradiometer (MODIS) Land Cover (MCD12Q1) version 6.1 data product (Friedl & Sulla-Menashe, 2022), accessed using the ‘modisfast’ package in R (Taconet & Moiroux, 2024) and calculated the dominant land cover type at each checklist location using the University of Maryland (land cover type 2) classification system (Hansen et al., 2000). We obtained coordinates of past wildfire boundaries from the National Burned Area Boundaries Dataset published by the Monitoring Trends in Burn Severity program (Eidenshink et al., 2007) and removed checklists in which more than 33% percent of a 1.5km radius circle surrounding the checklist location was affected by fire in the past 4 years. Measurements of snow depth on April 1 were obtained from the Snow Data Assimilation System (SNODAS) Data Product (National Operational Hydrologic Remote Sensing Center) and hourly precipitation and windspeed data were obtained from the European Centre for Medium-Range Weather Forecasts Reanalysis v5 product (Copernicus Climate Change Service, 2023; Hersbach et al., 2018). See the Supplemental Methods for more information on how environmental data were processed.

### Ensemble average approach for counts of mountain chickadees

Our restricted geographic, elevational, and date boundaries, combined with the relative remoteness of our focal area meant our sample sizes of eBird checklists were small compared to many previous studies of eBird data (e.g. Fink et al., 2020; Youngflesh et al., 2021). Therefore, instead of using training and testing methods (e.g. Strimas-Mackey et al., 2023), we used an ensemble modeling approach to predict mountain chickadee abundance, similar to the basic framework of Fink et al. (2020). After filtering for an elevation range, geographic region, and time frame, we first built a 3km x 3km grid over our region of interest, then randomly selected one eBird checklist per week, per grid cell. This process was repeated 100 times to create 100 sampling datasets for each elevation range within a region. For each sampling dataset, we modeled the relationship between the number of chickadees observed in each checklist (response variable) and multiple effort, time, and environmental variables (described below) using a generalized linear model with the negative binomial distribution (nbinom2 in the R package glmmTMB; Brooks et al., 2017; Hardin & Hilbe, 2007). For each of the 100 models, we generated predictions for the variable of interest (i.e. year or snow depth), then averaged the model predictions at each value on the x-axis (year or 1cm increments for snow depth) to obtain an ensemble average prediction for each elevation range within a region. This approach of selecting one checklist per grid cell per week allowed us to limit the spatial-temporal bias that is common in participatory science data (Johnston et al., 2021; Strimas-Mackey et al., 2023) and when combined with an ensemble framework, allowed us to use our entire dataset to predict mountain chickadee abundance.

### eBird model structure

Each model included multiple effort and time variables that are known to affect detection and abundance measures in eBird data (Johnston et al., 2021). To control for observer effort, we included checklist length (effort hours) and checklist distance (effort distance) in km as linear fixed effects. To control for time, we included year, day of year, and hour of day as linear fixed effects. It was unclear whether every environmental variable we sourced (i.e. land cover, latitude, total precipitation, and windspeed) would be an important predictor of mountain chickadee abundance. We tested the importance of each environmental variable using likelihood ratio tests that compared a baseline model with only the effort and time variables to models that also included one of the environmental variables as a fixed effect (see Supplemental Methods for more information). Our final model included all effort and time variables (checklist length, checklist distance, year, day of year, and hour of day) and dominant land cover class and latitude as fixed effects.

### Validation step 1: Measuring reproduction using eBird data

eBird data have been used to estimate avian reproduction in the past (e.g. Socolar et al., 2025), but to our knowledge, eBird observations have never been validated as a measure of breeding productivity by using on-the-ground measurements of individual breeding attempts. Our long-term study offered a unique opportunity to validate local eBird data as a measure of breeding productivity by comparing yearly variation in mean brood sizes at the long-term site to yearly variation in post-breeding counts of mountain chickadees in eBird data from the region surrounding our study site. However, it was not initially clear how large the geographic regions should be or what time frame of observations would best reflect population-level reproduction. Larger geographic regions would increase the sample sizes of checklists, possibly improving the accuracy of our predictions, but larger regions might also increase the likelihood that birds in the sample were experiencing different environmental conditions, which may obscure biological trends. Similarly, longer time frames could also result in larger sample sizes, but a longer time frame might increase the likelihood that chickadee abundance could be affected by post-fledging juvenile mortality. We accounted for this uncertainty by using an exploratory analysis to determine which combinations of geographic regions and time frames used to filter the eBird data resulted in a dataset of post-breeding chickadee counts (chickadees per km) that best correlated with yearly brood sizes in the long-term data (see Supplemental Methods). The geographic region and time frame combinations that produced the strongest correlation at each elevation are presented in the results.

### Validation step 2: Snow depth and reproductive trends in local eBird data

The first validation step revealed that post-breeding chickadee abundance calculated from eBird data can provide accurate estimates of yearly reproduction in mountain chickadees, but it was important to test whether breeding production measured via eBird data exhibited similar relationships to snow depth as observed in the long-term study. Using the geographic regions and time frames that best matched the yearly variation in reproduction at the long-term site, we applied our ensemble modeling approach to examine the relationships between chickadee counts in eBird data and snow depth within each elevation range to see if we observed similar relationships as seen in the long-term data. We employed the same model structure as described above, but here we included snow depth as a fixed effect instead of year and tested for a threshold effect of snow depth in each sampling dataset in the ensemble. Predictions from the model with the best-supported threshold value from each dataset, or the linear model if it had the lowest AIC value, were used to calculate the ensemble average. Ensemble averages built from models with a significant snow depth term (pre- or post-threshold) better explained the data (lower RMSE) than ensemble averages based on all models, so in later steps we calculated ensemble averages from models with a significant snow depth term (Figure S8). Due to some models being dropped for not having significant snow depth terms, here and in following steps we built ensemble model predictions from 200 sampling datasets instead of 100.

To confirm the findings our ensemble average approach, we also analyzed the relationship between snow depth and chickadee counts in eBird data using single generalized linear mixed models that included site ID as a random effect. We explored the robustness of the relationships between snow depth and post-breeding counts of chickadees by examining how these relationships changed with adjustments to the geographic regions and time frames used for filtering the eBird data (see Supplemental Methods).

### Exploration of snow depth and reproduction relationships across the Sierra Nevada

Our validation analyses provided confidence that 1) post-breeding eBird counts of mountain chickadees can reflect yearly variation in brood sizes observed in the long-term study (Figure 2), 2) relationships between snow depth and reproduction observed in the long-term study can be reproduced using eBird data and SNODAS model predictions of snow depth from the region surrounding the long-term study (Figure S8, S9), and 3) relationships between snow depth and reproduction are fairly robust to changes in the geographic limits and time frames used to filter the eBird data (Figure S10, S11). This 3^rd^ finding was especially important for exploring relationships between snow depth and chickadee reproduction in new regions where we did not have long-term data to validate the filters. The apparent robustness of these relationships gave us confidence that we could apply the same methods to regions beyond our long-term study site.

**Figure 2.**
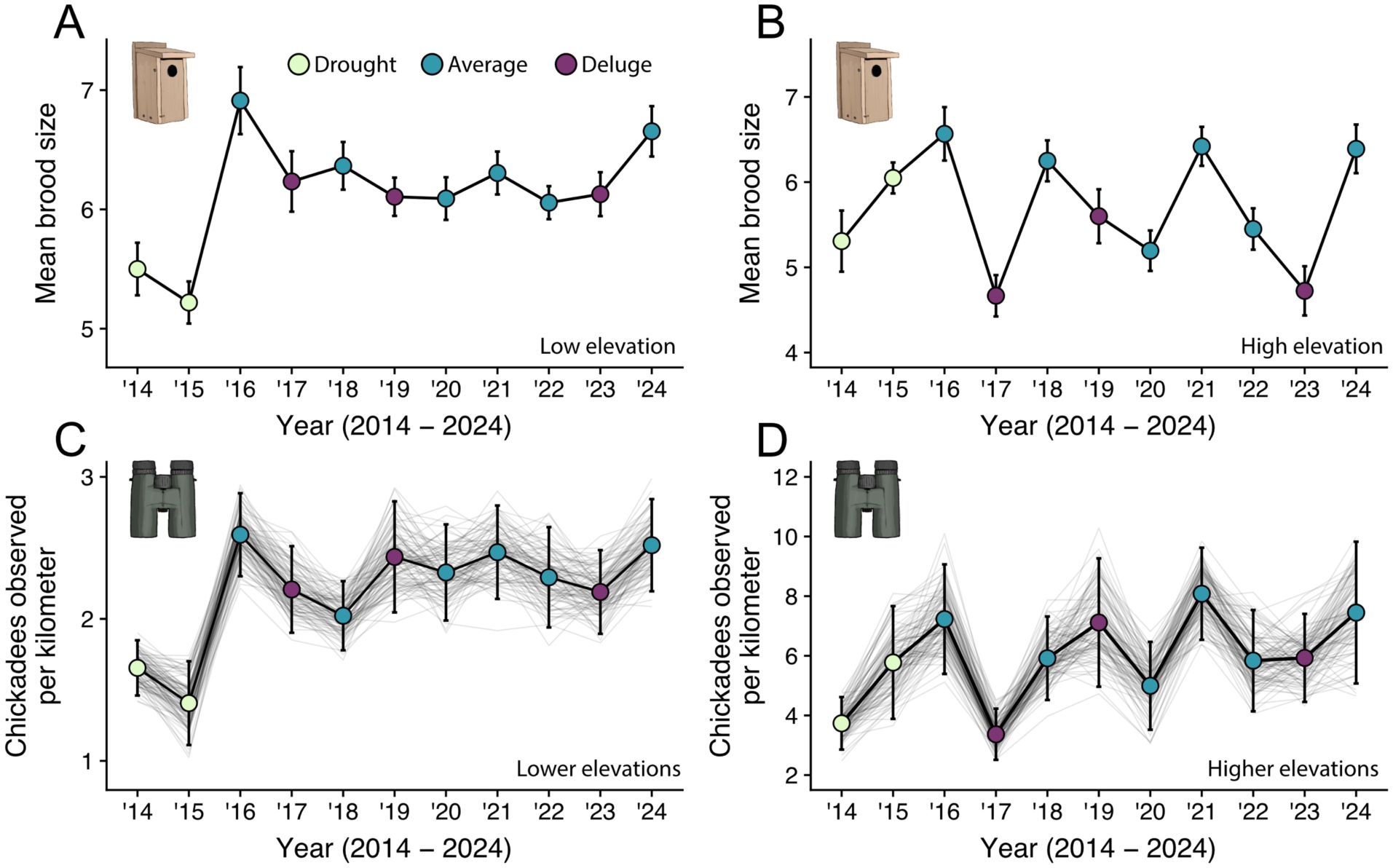
Comparison of yearly variation in mountain chickadee brood sizes at the long-term study site to yearly variation in post-breeding counts of chickadees in eBird data from the region surrounding the study site. Points represent data; connecting lines are presented as visual aids. A) Average (± SE) brood sizes by year at the low elevation site in the long-term study. B) Average (± SE) brood sizes by year at the high elevation site in the long-term study. C) Yearly variation in post-breeding chickadee counts in eBird data from lower elevations, corresponding to the low-elevation site in the long-term study (1829-2134m; 6000-7000ft). D) Yearly variation in post-breeding chickadee counts in eBird data from higher elevations, corresponding to the high-elevation site in the long-term study (>2377m; >7800ft). Light gray lines in C and D connect individual model predictions (points not shown). Black connecting lines and colored points in C and D show the ensemble averages of the predictions (mean of the individual models). Error bars in C and D show 95% reference intervals. Data from Cornell Lab | eBird. Points are colored relative to yearly snowfall (drought conditions, average conditions, deluge conditions).

We explored how relationships between snow depth and reproduction varied across the Sierra Nevada mountains by applying our ensemble average modeling approach to overlapping latitudinal windows. As above, each geographic range was constrained by the east-west boundaries of our perimeter, but in this step, we standardized the north-south extents of the latitude windows within each elevation range at extents that maximized sample sizes but limited the potential of the window to encompass different habitats. At lower elevations we used windows that extended 1.2° north-south in latitude, and at upper elevations we used windows that extended 1.6° north-south in latitude due to fewer eBird checklists being submitted from higher elevations. Latitude windows shifted by 0.4° at both elevations with each increment. This overlapping approach allowed us to observe shifting relationships between snow depth and reproduction across regions with different snow regimes.

For time frames, we used the same post-breeding time ranges in this broader analysis as those previously identified using the eBird data near the long-term study site (see Results). It is possible that populations at southern latitudes could finish breeding earlier than populations at northern latitudes, but the few published records of timing of breeding from mountain chickadee populations in the Sierra Nevada suggest that timing of breeding is fairly consistent across latitudes (see Supplemental Methods). Ensemble averages were calculated from sampling runs in which the pre- or post-threshold slope term was a significant predictor of chickadees observed (P<0.05).

## Results

### Snow depth and reproduction in the long-term study

Yearly snow depth varied substantially in the long-term study, but the high elevation site consistently received more snowfall each winter than the low elevation site (Figure 1A). Winter snow accumulation affected offspring production in the following breeding season, but the effect of snow depth on mountain chickadee reproduction differed across elevations, matching previous findings from this population. At lower elevations (1965m – 2070m), offspring production was negatively affected by low-snow winters: chickadees produced on average 1 fewer offspring in drought years versus average and deluge years (N=543, β=-0.16, χ^2^1=36.48, P<0.001). Modeling snow depth on a continuous scale revealed that brood sizes at lower elevations increased sharply between 0 and 38cm of April 1 snow depth (β=0.006, χ^2^1=31.10, P<0.001), then remained stable following April 1 snow depths between 38cm and 94cm (β=-0.0001, χ^2^1=0.67, P=0.413, Figure 1B). In contrast, reproduction at high elevations (2380m – 2590m) was negatively affected by high-snow winters: chickadees produced on average 1 fewer offspring in deluge years versus drought and average years (N=386, β=-0.16, χ^2^1=21.37, P<0.001). On a continuous scale, brood sizes at higher elevations were relatively stable following 89cm to 367cm of April 1 snow depth (β=0.0004, χ^2^1=2.10, P=0.148), but brood sizes declined when April 1 snow depth exceeded 367cm (β=-0.003, χ^2^1=13.91, P<0.001, Figure 1C).

### Using eBird data to measure yearly reproduction

Our exploratory analyses revealed that model predictions based on post-breeding counts of mountain chickadees in eBird data can reflect yearly variation in offspring production at the long-term study site (validation step 1). At lower elevations, model predictions of chickadee counts based on eBird data collected from July 17 to Sept 9 (end of the 36^th^ week of the year) and between latitudes 39.2° to 39.6° N correlated strongly with yearly variation in brood sizes at the low elevation site in the long-term study (N=1586 checklists, P<0.001, Adj. R^2^=0.77, Figure 2, S3). At higher elevations, the best-matching time frame in the eBird data was shorter and the geographic region was larger, the latter likely due to the smaller number of checklists submitted from high elevations. Model predictions based on eBird data collected from July 29 to Aug 19 (end of the 33^rd^ week of the year) and between 38.6° to 40.2° N at higher elevations correlated with yearly variation in brood sizes at the high elevation site in the long-term study (N=452 checklists, P=0.007, Adj. R^2^=0.53, Figure 2, S3). Importantly, many additional combinations of time frames and geographic regions centered on the long-term study site correlated with the long-term data at both elevations (P<0.05, Figure S4, S6), including the latitudinal extents used in the exploratory analyses below.

Modeling eBird data and snow depth near the long-term study site (validation step 2) revealed similar relationships between snow depth and reproduction as the long-term study. At lower elevations, the number of chickadees observed per kilometer during the post-breeding period increased below a low snow depth threshold (mode=23cm and mean=62cm using the ensemble approach, and 37cm using a single model with checklist location as a random effect), then stabilized after the threshold (Figure S8, S9), similar to the long-term study. At higher elevations, the number of chickadees observed per kilometer showed a slight positive slope below the snow depth threshold (mode=357cm and mean=315cm using the ensemble approach, and 357cm using a single model with checklist location as a random effect), then showed a steep decline beyond the threshold (Figure S8, S9), also similar to the long-term study. Importantly, relationships between snow depth and chickadee reproduction at both elevations were consistent across time frames and geographic regions used to filter the eBird data, suggesting these relationships are robust to minor differences in the time and geographic parameters used (Figure S10, S11).

### Snow depth and reproduction across the Sierra Nevada

Mountain chickadee populations across the Sierra Nevada experienced vastly different snowfall regimes depending on their elevation and latitude. Lower elevations consistently received less snow than higher elevations at the same latitudes, and latitude itself was strongly associated with snowfall, with northern latitudes receiving more snow that lasted longer into the summer than southern latitudes (Figure 3). Snow depth influenced chickadee reproduction at both lower and higher elevations across the Sierras, but the relationship between snow depth and reproduction changed across elevations and latitudes.

**Figure 3.**
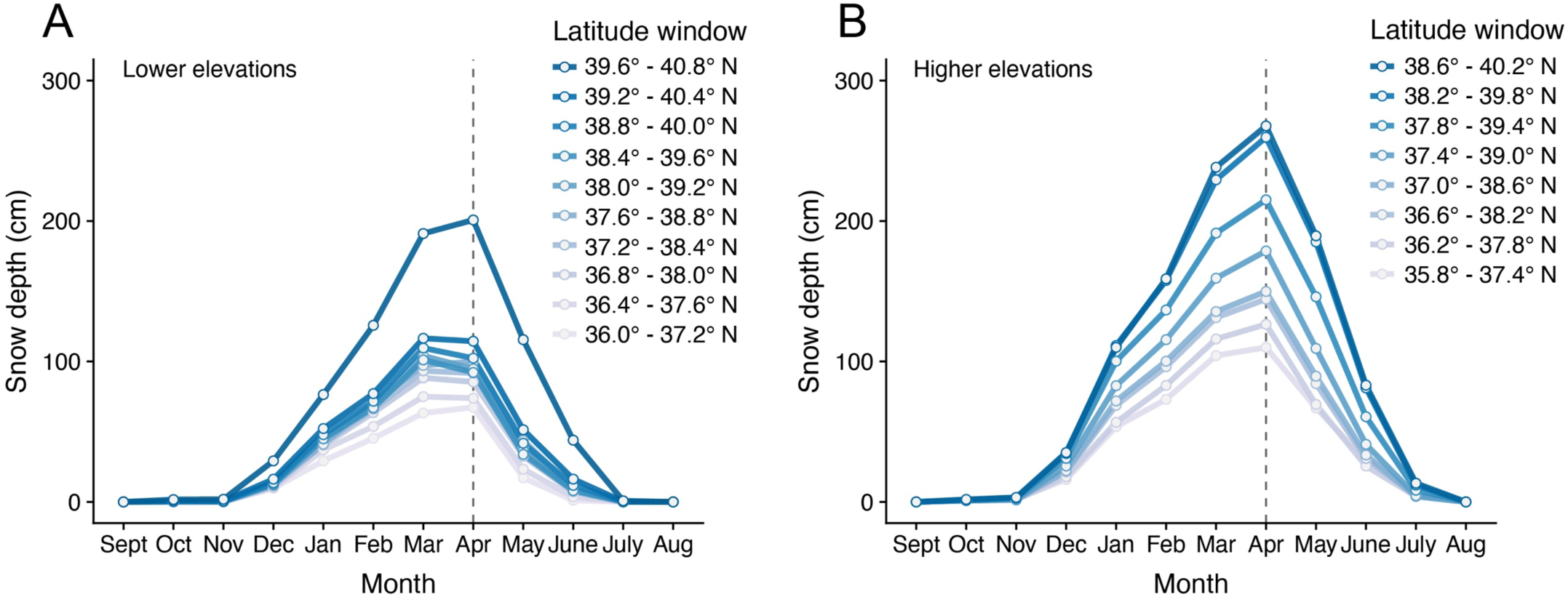
Snow depth is variable across latitudes and elevations in the Sierra Nevada. Average monthly snow depth at eBird checklist locations by latitude window from 2014 to 2024 for A) elevations between 1829 and 2134m (6000-7000ft), corresponding to the low-elevation site in the long-term study, and B) elevations equal to or above 2377m (7800ft), corresponding to the high-elevation site in the long-term study.

Our exploratory analysis revealed that chickadee reproduction at lower elevations across the Sierras was mostly unaffected by high snow conditions, as in the long-term study. However, low elevation populations within the most northerly latitude window (39.6° - 40.8° N; north of the long-term study) exhibited a negative relationship between reproduction and snow depth (Figure 4C), possibly a result of these populations experiencing much higher snow depths than the populations to the south (Figure 3A, uppermost line vs all other lines). Low elevation populations within the latitude windows that included the long-term study site exhibited very similar relationships to snow depth as to what we observed in the long-term study. Reproduction in these areas was lower during drought conditions, then remained fairly stable after achieving a relatively low snow depth (Figure 4D-F). Median threshold values for these three windows ranged from 54cm snow depth (Figure 4D) to 5cm snow depth (Figure 4E and 4F). Moving further south, low elevation populations within the next two latitude windows appear to show a beneficial effect of higher April 1 snow depth levels, with reproductive output increasing slightly following higher snow conditions (Figure 4G, 4H). The following three windows to the south showed weak, if any relationship between chickadee reproduction and snow depth (Figure 4I-K), and the final latitude window shows a general negative effect of snow on reproduction, but the individual models used to build the ensemble average did not agree on a common threshold value or relationship so this result should be interpreted with caution (Figure 4L).

**Figure 4.**
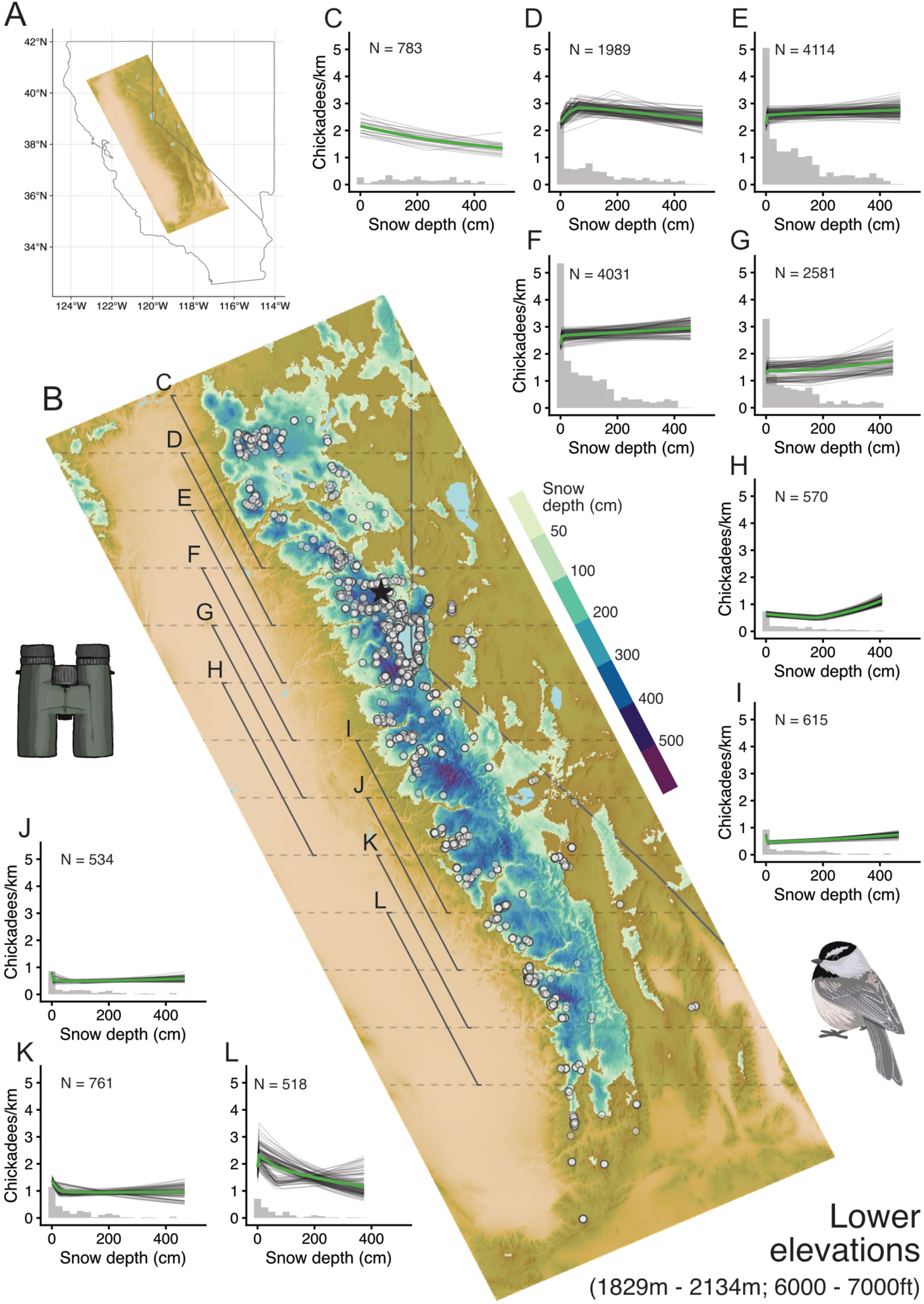
Relationships between snow depth and mountain chickadee reproduction are variable across the Sierra Nevada at lower elevations (1829m-2134m; 6000-7000ft), corresponding to the low elevation site in the long-term study. A) Map key. B) Map of snow depth on April 1^st^, 2019 (a deluge year) with the locations of eBird checklists from 2014 to 2024 collected at elevations between 1829m and 2134m (white circles); data from Cornell Lab | eBird. Black star shows the location of the long-term study site. Dashed gray lines show latitude lines every 0.4° and solid diagonal gray lines show the range of eBird and snow data included in each latitude window corresponding to the model prediction plots. C-L) Plots C through L show the relationship between snow depth on April 1^st^ and post-breeding counts of mountain chickadees in eBird data within each latitude window. Gray lines in C-L show individual model predictions and green lines show the ensemble average of the model predictions. Gray histograms show the number of eBird checklists associated with 25cm increments of snow depth. Uncertainty in the model predictions are often associated with low sample sizes. A value of 1 on the y-axis equals 250 eBird checklists. N values in each plot show the sample size of eBird checklists included within each latitude window: C) 39.6° - 40.8° N, D) 39.2° - 40.4° N, E) 38.8° - 40° N, F) 38.4° - 39.6° N, G) 38° - 39.2° N, H) 37.6° - 38.8° N, I) 37.2° - 38.4° N, J) 36.8° - 38° N, K) 36.4° - 37.6° N, L) 36° - 37.2° N.

As in the long-term study, the high elevation populations within the latitude windows that included the long-term study site showed a strong negative effect of high-snow conditions on reproduction, but in contrast to the long-term study, these populations also showed a more positive slope before the threshold snow depth, suggesting the presence of an optimal mid-level snow depth for reproduction at higher elevations (Figure 5C-E). Median snow depth thresholds for these three regions ranged from 299cm to 317cm snow depth. To the south, in the central Sierra Nevada, the relationship between snow depth and reproduction changed dramatically.

**Figure 5.**
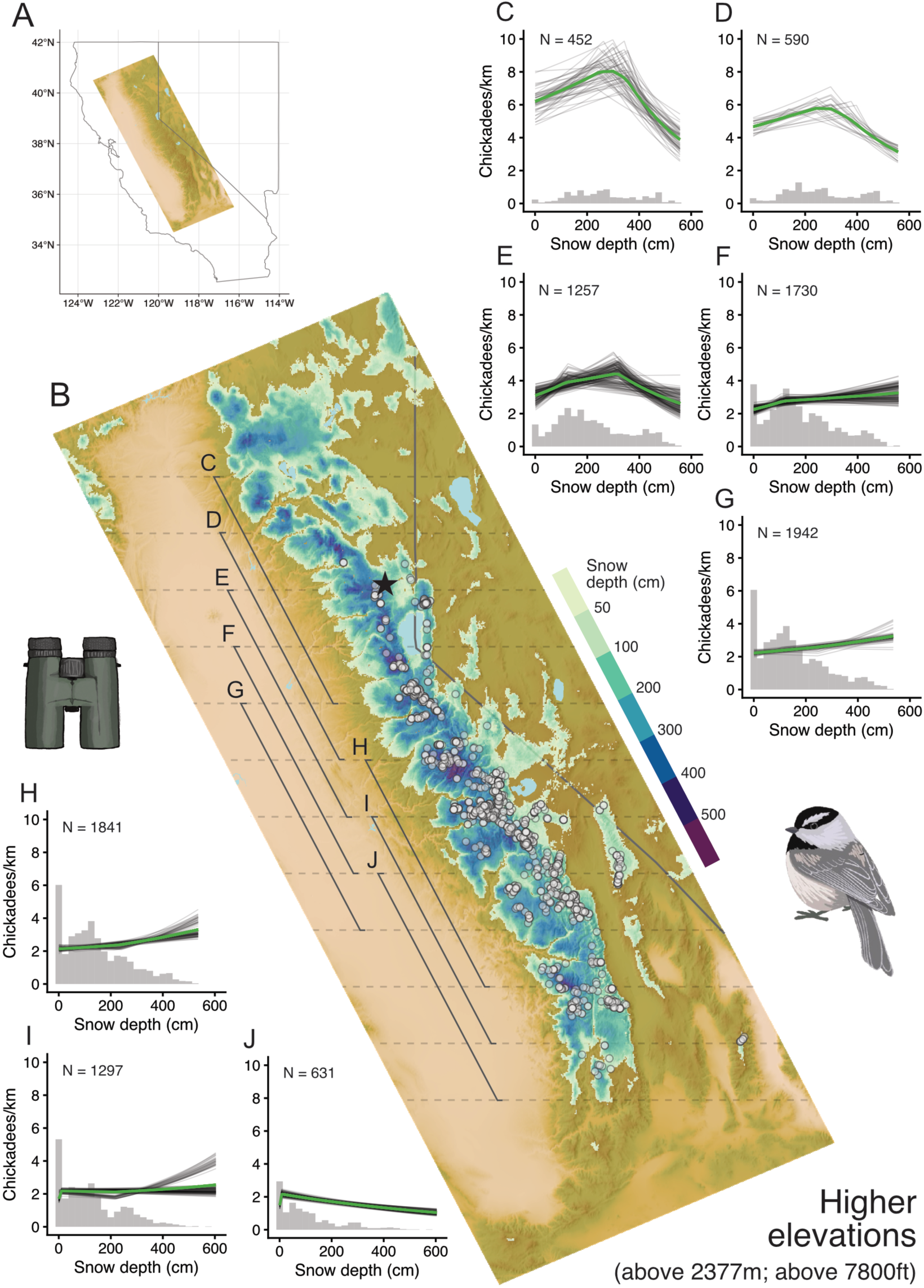
Relationships between snow depth and mountain chickadee reproduction are variable across the Sierra Nevada at higher elevations (equal to or above 2377m, 7800ft), corresponding to the high elevation site in the long-term study. A) Map key. B) Map of snow depth on April 1^st^, 2019 (a deluge year) with the locations of eBird checklists from 2014 to 2024 collected at elevations above 2377m (white circles); data from Cornell Lab | eBird. Black star shows the location of the long-term study site. Dashed gray lines show latitude lines every 0.4° and the solid diagonal gray lines show the range of eBird data and snow depth data included in each latitude window, corresponding to the model prediction plots. C-J) Plots C through J show the relationship between snow depth on April 1^st^ and post-breeding counts of mountain chickadees in eBird data within each latitude window. Gray lines in C-J show individual model predictions and green lines show the ensemble average of the model predictions. Gray histograms show the relative number of eBird checklists associated with 25cm increments of snow depth. Uncertainty in the model predictions are often associated with low sample sizes. A value of 2 on the y-axis corresponds to a count of 100 eBird checklists. N values in each plot show the sample size of eBird checklists included within each latitude window. C) 38.6° - 40.2° N, D) 38.2° - 39.8° N, E) 37.8° - 39.4° N, F) 37.4° - 39° N, G) 37° - 38.6° N, H) 36.6° - 38.2° N, I) 36.2° - 37.8° N, J) 35.8° - 37.4° N.

Populations in the three central latitude windows showed a beneficial effect of snow depth on reproduction, with more chickadees being observed in the post-breeding period following high-snow winters compared to low-snow winters (Figure 5F-H). In contrast, snow depth appeared less important for reproduction at higher elevations in the southern Sierras, with the second-to-last latitude window showing slight negative effect of drought conditions (Figure 5I), and the final latitude window showing very little effect of snow depth on reproduction (Figure 5J).

## Discussion

Extreme climatic events are increasing in frequency, but the impact of these events on animal populations will likely vary across environments. By combining our long-term study of mountain chickadees with participatory science observations, here we reveal that drought and deluge snow extremes generate differential reproductive responses across elevations and latitudes, adding to growing recognition that biological responses to environmental conditions are often nonstationary, or variable across spatial scales (Lindenmayer et al., 2024; Rollinson et al., 2021). Our study also shows how insights from long-term studies can be used to inform broader analyses of participatory science data investigating how reproduction within populations inhabiting different environments is differentially affected by environmental extremes.

### Snow depth and reproduction in the Northern Sierra Nevada

Drought and deluge snowfall extremes are increasingly common in the western United States (Huning & AghaKouchak, 2020; Swain et al., 2018), but these extremes result in variable snow depth patterns across elevations and latitudes (Huning & AghaKouchak, 2018). Our long-term study of mountain chickadees has revealed that reproductive performance is strongly associated with spring snow depth in the northern Sierra Nevada mountains. At lower elevations, the effect of spring snow depth on reproduction was mostly determined by its presence or absence: only 38cm (approximately 15 inches) of snow depth on April 1 was required for average reproductive performance at the low elevation site and exploratory analyses of eBird data confirmed this conclusion while suggesting this threshold could be even lower in some regions (5cm to 54cm; Figure 4D-F). High spring snow depths did not affect reproduction at low elevations in the long-term study, but April 1 snow depth never exceeded 94cm at the low elevation site. Some low elevation populations in the latitude windows that included our long-term site did experience high spring snow depths (Figure 4D-F histograms), but the effect of high snow depth on low elevation reproduction in the eBird data was mixed. High spring snow depths in the southern two latitude windows that included our long-term site had no effect on reproduction (Figure 4E and 4F), but reproductive output did decline slightly with increasing snow depth in the northern low-elevation latitude window that included our long-term site (Figure 4D), suggesting that northern populations may be negatively affected by high snow depths even at low elevations.

At higher elevations, we observed no negative effects of low snow depths on reproduction in the long-term study, but the exploratory analyses of eBird data challenged this conclusion. April 1 snow depth was never lower than 88cm at the high elevation site in the long-term study, but some of the high elevation populations in the eBird analyses experienced much lower spring snow depths. The three high elevation latitude windows that included the long-term study site showed very similar threshold values as the long-term study, but their pre-threshold slopes were much steeper than the long-term study, suggesting reproduction in these populations was negatively affected by low spring snow depths (Figure 5C-E). However, in both the long-term study and the eBird analyses, the effect of low snow depth on reproduction at high elevations was far outweighed by the negative effect of high snow depths. More than 367cm (144 inches, or 12 feet) of snow depth on April 1 was associated with a sharp decline in reproductive output at the high elevation long-term site and this was confirmed in the eBird analyses that showed similarly sharp declines after median snow depth thresholds ranging from 299cm to 317cm (Figure 5C-E).

Previous investigations from our long-term study indicate that differential responses to snow depth across elevations may be a result of both local adaptations and universal responses to constraints on reproduction imposed by environmental extremes. The close proximity of our low and high elevation sites (3.5km) may appear to preclude selection for local adaptation, but intraspecific local adaptation across elevational gradients has been observed in birds (Atwell et al., 2014; McCormack & Smith, 2008; Robertson et al., 2026) and other animal taxa (Bachmann et al., 2020). Indeed, chickadees living at higher elevations exhibit a number of adaptations to the harsher, longer winters at higher elevations compared to the milder, shorter winters at lower elevations. High elevation mountain chickadees exhibit enhanced spatial cognition and larger hippocampi with more hippocampal neurons compared to low elevation birds (Croston et al., 2016; Freas et al., 2012), whereas low elevation chickadees are bolder, more aggressive, exhibit higher cognitive flexibility, and sing different songs than high elevation chickadees (Branch et al., 2015; Croston et al., 2017; Kozlovsky et al., 2014, 2015). Further, chickadees living at our higher elevation study site produce fewer, heavier offspring than chickadees at lower elevations who produce more, lighter offspring (Sonnenberg et al., 2025). Elevational variation in nestling mass is commonly observed across elevational gradients in birds (Bears et al., 2009; Boyle et al., 2016) and this trait is positively associated with recruitment rates in this species (Whitenack et al., 2025). These differences suggest that local adaptation can occur across elevations separated even by short distances and in the presence of gene flow (Branch et al., 2017); thus, some of the reproductive responses to snow depth we observed are likely associated with local adaptation to different snow regimes. For example, chickadee reproductive output at lower elevations was fairly stable following April 1 snow depths between 50cm and 250cm, but high elevation April 1 snow depths within this range resulted in slightly reduced reproductive output in the eBird data,

suggesting high-elevation birds may be less adapted to low-snow conditions than low-elevation birds.

Alternatively, drought and deluge snowfall regimes may be so extreme that they impose universal constraints on avian reproduction that would affect chickadees breeding at any elevation. Insights into the local environmental predictors of timing of breeding and reproductive output from our long-term study provide evidence for this hypothesis. At low elevations, timing of breeding and clutch sizes are not associated with snow depth, but mountain chickadee brood sizes are positively associated with autumn snow accumulation (Whitenack et al., 2023). This suggests that low elevation chickadees begin breeding at ‘normal’ times in drought years and lay similar clutch sizes to those laid in average and high snow years, but drought conditions result in fewer offspring surviving to the fledging stage. The most likely mechanism for this decline in nestling survival is a negative effect of drought conditions on insect abundance (Forister et al., 2018; Halsch et al., 2024). For example, many invertebrates spend the winter underground protected by an insulative layer of snow, but low winter snow cover can increase overwinter mortality when insects are exposed to freezing temperatures (Bale & Hayward 2010). Increased insect mortality will result in reduced food availability and reduced nestling survival in songbirds like chickadees that rely on insects to feed their young (Grames et al., 2023; Lack, 1968).

At high elevations, the extreme snow depths and late snowmelt following deluge conditions delayed initiation of breeding in mountain chickadees in our long-term study (Whitenack et al., 2023) and this same pattern is seen in other high elevation and high latitude breeding species (Brawn et al., 1991; Bison et al., 2020; Hendricks, 2003). Late initiation of breeding is commonly associated with reduced clutch sizes in northern-hemisphere breeding songbirds (Goodenough et al., 2009, Perrins & McCleery, 1989), and correspondingly, first egg dates are negatively correlated with clutch and brood sizes in mountain chickadees (Whitenack et al., 2024, 2025). Thus, in deluge years, long-lasting snow cover results in later breeding, smaller clutch and brood sizes, and fewer offspring fledged. The exact mechanism for why snow cover inhibits breeding is unclear, but long-lasting snow may cover nesting material, obscure cavity entrances, or delay insect emergence (Whitenack et al., 2023).

### Snow depth and reproduction across the Sierra Nevada

Exploration of eBird data from across the Sierra Nevada mountains revealed the trends we observed in the long-term study were somewhat localized to the northern Sierras. Populations to the north and south of the long-term study exhibited different relationships between spring snow depths and reproduction compared to the long-term study, and populations to the south also showed remarkably less variation in their responses across elevations. The average reproductive trend at both low and high elevations in the central Sierras was that higher spring snow depths increased reproductive output (Figure 4G-I, Figure 5F-H). This is a dramatic shift from the populations near our long-term field site where high snow depths clearly had a negative effect on reproduction at high elevations and often had no impact at low elevations. The most likely cause of this difference is the timing of snow melt. Snow melted earlier in the central Sierra than the northern latitudes (Figure 3), meaning extreme snow depths following deluge conditions in the central latitudes may not delay initiation of breeding if snow melts early in the spring. Instead, the positive relationship between snow depth and reproduction in the central Sierra suggests that increased spring snow depth may result in increased insect food availability (e.g., via increasing water availability into the summer or improving overwinter survival of insect populations), leading to more nestlings surviving to the fledging stage and therefore higher reproductive output. Low elevation populations in the central Sierras were also less affected by drought conditions compared to the northern Sierras, suggesting these populations may be better adapted to low-snow conditions, or that central Sierra populations rely on an insect food base that is better adapted to low-snow conditions. Additional long-term studies that pair insect abundance data with data on vertebrate reproduction will be key to unpacking these hypotheses.

Snow depth had little effect on reproduction at low elevations in the southern Sierras where spring snow depths were lowest (Figure 3) and ensemble model predictions were mixed, possibly due to lower sample sizes of eBird checklists from these regions (Figure 4K and L).

However, high elevation populations in the southern Sierras showed relationships that resembled those exhibited by the low elevation populations in the northern Sierras in which drought conditions reduced reproductive output, but higher snow depths were associated with mostly stable (Figure 5I) or declining reproductive performance (Figure 5J). Winter temperatures are warmer at these southern latitudes and winter precipitation is more likely to occur as rain (Lundquist et al., 2008). In such regions it might not be surprising that snow depth has less effect on reproduction, especially at lower elevations.

These findings reveal substantial variation in mountain chickadee reproductive responses to spring snow depths across the Sierra Nevada range. Deluge snow conditions decreased reproductive output in northern, high elevation mountain chickadee populations, but improved reproductive performance at both low and high elevations in the central Sierra Nevada. Drought conditions decreased reproduction at low elevations in the northern Sierra and at both low and high elevations in the central Sierra. Such differential responses across populations complicate ecological forecasts, species distribution modeling, and conservation planning, making it difficult to predict future population trajectories (Pease et al., 2022a; Rollinson et al., 2021).

These differential responses may appear to suggest that mountain chickadees will have sources of population stability despite increasing extreme climate events. For example, even when high elevation populations at northern latitudes produced fewer young following deluge conditions, low elevations produced average numbers of young, meaning low elevation populations could potentially act as source populations for high elevations following low reproductive performance in deluge years. However, the extreme degree of elevational adaptation this species exhibits (e.g. Pravosudov et al., 2025) could limit this process. Low elevation populations experience milder winters and less selection for cognitive abilities to remember the locations of cached food (Croston et al., 2016), meaning individuals dispersing from low elevations may be less likely to survive at high elevations where winters are longer and harsher. Further investigations of the interactions between local adaptation, reproduction, and selection across different environments will be required to better address these hypotheses.

## Conclusions

Our study reveals that mountain chickadee reproductive responses to snowfall extremes differ across elevations and latitudes in the Sierra Nevada, likely due to adaptation to local environments as well as universal responses to environmental constraints imposed on reproduction by drought and deluge snow extremes. Increasing evidence suggests that spatial nonstationarity of biological responses to environmental variation is common in vertebrate taxa (Evans et al., 2024; Johnson et al., 2023; Pease et al., 2022b; Socolar et al., 2025; Tonelli et al., 2024; Youngflesh et al., 2021), but accurately measuring variation in reproduction across environments remains a major challenge for understanding animal responses to environmental change. Long-term studies should continue to lead this work due to the invaluable insights that can be obtained by studying marked populations, including identification of the individual-level mechanisms that underly population-level trends. However, despite the importance and irreplaceability of long-term studies in a changing world, they are sometimes limited in their scope and application. For example, if our long-term research had been conducted in the central Sierra instead of the northern Sierra, we would have come to nearly the opposite conclusion about the effects of snow depth on reproduction at high elevations. Yet, this limitation cannot be solved entirely by participatory science observations collected across a broader scale.

Participatory science observations are transforming our understanding of animal responses to environmental change (Evans et al., 2024; Senzaki et al., 2020; Socolar et al., 2025), but they often lack the nuanced view into individual responses to environmental variation provided by long-term studies. Without validating participatory science data as an accurate measure of complex processes like reproduction, conclusions based on such data may be misleading.

Our findings suggest that using long-term data to validate, then interpret participatory science data offers a powerful framework for studying how differences in individual reproductive responses to environmental variation lead to spatial nonstationarity in population-level demography. We encourage the leaders of other long-term studies to explore whether this framework or a similar framework can be used in their systems to further our understanding of how biological responses to environmental variation change across major geographic scales and allow us to better predict future population-level responses to environmental change.

## Acknowledgements

We would like to thank Drs. Rebecca Croston, Maria Tello-Ramos, Dovid Kozlovsky, Lauren Benedict, Virginia Heinen, and Yuting ‘Hermione’ Deng for assistance with data collection. Jeff Brown and Dan Sayer of Sagehen Creek Field Station (University of California Berkeley) provided assistance at our field site. We also thank the many birders who submitted the eBird checklists that made this project possible. Nest box and binocular drawings by Sofia Haley and chickadee drawings by Katherine Johnson.

