## Supplemental material for "Drought to deluge: Differential impacts of snow on mountain chickadee reproduction across the Sierra Nevada mountains"

### Supplemental methods

*Snow depth in the long-term study*

The Independence Lake SNOTEL (site 541, elevation 2541m, coordinates: 39.4° N, −120.31° W) is located within the high-elevation study site, and the Independence Camp (site 539, elevation 2128m, coordinates: 39.45° N, −120.29° W) and Independence Creek (site 540, elevation 1962m, coordinates: 39.49° N, −120.28° W) SNOTEL sites are located near the low-elevation study site (4.5km and 6km away) in the same watershed and along the same exposure. As a result, the average snow depth of the Independence Camp and Independence Creek sites accurately represents the mean snow depth across the low-elevation site (Kozlovsky et al. 2018, Whitenack et al. 2023).

*Environmental data*

We obtained information on environmental conditions in the Sierra Nevada mountains from multiple sources to help predict mountain chickadee abundance in eBird data. The spatial resolution of the elevation data obtained from the Shuttle Radar Topography Mission (SRTM; NASA JPL) Mapzen terrain tiles (Tilezen 2025), courtesy of the U.S. Geological Survey is approximately 30m. The spatial resolution of the dataset is 500m MODIS Land Cover

(MCD12Q1) version 6.1 data product (Friedl and Sulla-Menashe 2022) is 500m. Land cover data for 2024 was not available when we accessed these data, so for checklists collected in 2024, we used the 2023 land cover classifications. Our dataset of eBird checklists was relatively small due to our focus on limited geographic and time ranges, so unlike previous studies with much larger datasets, we did not calculate predictors for each land cover type (e.g. Fink et al. 2020). Instead, we calculated the dominant land cover class within a 1.5km radius circle centered on each eBird checklist location and removed checklists where the dominant land cover was "urban" due to the unnatural habitat of those locations.

High-severity wildfires have reshaped many landscapes in the Sierra Nevada in recent years (Williams et al. 2023), so we removed eBird checklists from areas recently affected by fire, where chickadee abundance had likely changed (Ray et al. 2025). The spatial resolution of the National Burned Area Boundaries Dataset is 30m (Eidenshink et al. 2007).

Snow depth data from the Snow Data Assimilation System (SNODAS) Data Product (National Operational Hydrologic Remote Sensing Center) come from model predictions (point estimates) of snow depth across the United States at a 1km resolution. Comparing SNODAS and SNOTEL measurements at the SNOTEL sites near the long-term study site revealed that the SNODAS model predictions are extremely highly correlated with the point measurements at SNOTEL sites (adjusted r-squared values > 0.92, supplemental figure S2), suggesting that SNODAS data are accurate predictions of snow depth in the Sierra Nevada mountains.

Detection of mountain chickadees could be affected by local weather conditions, so we obtained total hourly precipitation and windspeed data from the European Centre for Medium-Range Weather Forecasts Reanalysis v5 product (Copernicus Climate Change Service 2023, Hersbach et al. 2018). 10m windspeed u- and v-components were downloaded separately then combined to calculate wind velocity using the equation: velocity=sqrt(u^2 + v^2) (Hersbach et al. 2018). Hourly precipitation and windspeed data were matched to each eBird checklist using the hourly weather values closest to the middle time point of the checklist period. The resolution of these weather data is approximately 31km.

*eBird model structure*

We tested the importance of each environmental variable using a preliminary version of our final dataset that included eBird checklists collected before September 1 in each year. This end date was chosen to reduce the likelihood that predation and starvation of juveniles would impact count totals. Post-fledging mortality is high in songbirds (Naef-Daenzer and Grüebler 2016) and juvenile mortality is high in the first winter in mountain chickadees (Branch et al. 2019, Sonnenberg et al. 2019), so observations collected long after breeding may be less reflective of breeding production. We centered the geographic ranges for each elevation on the latitude of the long-term study site (approximately 39.4° N) and based the north-south extents of these ranges on early model validation runs. For lower elevations (1829-2134m, 6000-7000ft), we included checklists collected between 39.2° N and 39.6° N that were within the east-west bounds of the geographic perimeter described above. Far fewer checklists were submitted from high elevations (above 2377m, 7800ft), so we included checklists collected between 38.6° N and 40.2° N for higher elevations to obtain a large enough sample size for modeling. These end dates and geographic extents were further refined and validated in later steps.

We determined the importance of each environmental variable by incorporating likelihood ratio tests into our ensemble approach. Likelihood ratio tests can be used to compare a baseline model to a version of the same model that includes an additional fixed effect of interest to determine whether the addition of the new predictor improves upon the baseline model. Our implementation compared a baseline model with only the effort and time variables to a model that also included one of the environmental variables as a categorical (dominant land cover) or linear (latitude, total precipitation, wind velocity) fixed effect. We ran these tests for each of the 100 sampling datasets in the ensemble for each environmental variable and determined how often the environmental model improved upon the baseline model ($p<0.05$). The structure of the datasets used in each ensemble run was maintained for each environmental variable so that the importance of each variable was tested using the same data.

At lower elevations, dominant land cover improved upon the baseline model (likelihood ratio test $P<0.05$) in 39% of model runs, latitude improved 94% of model runs, and total precipitation improved 8% of model runs, whereas wind velocity did not improve any model runs. At high elevations near the long-term study site, dominant land cover improved the baseline model in 89% of model runs, latitude improved 100% of model runs, total precipitation did not improve any runs, and wind velocity improved 2% of model runs. Based on these results, we included only dominant land cover and latitude from these environmental variables in our final model structure.

*Validation step 1: Measuring reproduction using eBird data*

We used an exploratory analysis to find which combinations of geographic regions and time frames resulted in a dataset of post-breeding chickadee eBird counts that best correlated with average yearly brood sizes in the long-term data. As above, each geographic range increment was centered on the latitude of the long-term study site (39.4° N) and extended north and south in equal degrees. The smallest range increment extended from 39.3° N to 39.5° N and additional increments lengthened by 0.1° to the north and south to reach 38° N to 40.8° N at the largest range. Time frames started after the dates by which 99% of nests had fledged in the long-term study (described above), but we explored how well eBird datasets with different ending dates correlated with the long-term data. Ending dates for low elevations ranged from the end of the 31$^{st}$ week of the year (August 5$^{th}$ on a 365-day calendar) to the end of the 39$^{th}$ week of the year (September 30$^{th}$ on a 365-day calendar) and differed by 1-week periods. At high elevations, ending dates ranged from the end of the 32$^{nd}$ week of the year (August 12$^{th}$ on a 365-day calendar) to the end of the 39$^{th}$ week of the year due to fledging dates being later at higher elevations. These parameters resulted in 126 geographic range * time frame combinations at lower elevations and 112 combinations at higher elevations.

We compared yearly post-breeding mountain chickadee abundance (chickadees per km) in each of these eBird datasets to average yearly offspring production in the long-term data. For each eBird dataset associated with a geographic range * time frame combination, we used our ensemble modeling approach to calculate yearly post-breeding mountain chickadee abundance (chickadees per km) by including year as a fixed effect in each model. These models were run separately for each elevation range, then the model predictions for yearly abundance within each elevation range were correlated with the mean yearly brood sizes from the respective low or high elevation long-term study site using linear models (Figures S3-S6).

*Validation step 2: Snow depth and reproductive trends in local eBird data*

We examined the relationships between chickadee counts in eBird data and snow depth within each elevation range to see if we observed similar relationships as seen in the long-term data. When comparing support for snow depth threshold values here and in later steps, we considered snow depth thresholds from the 20% quantile to the 80% quantile of the snow depth term in each sampling dataset. If the 20% quantile was within 5cm of the lowest snow depth value in the sampling dataset, we examined support for threshold values starting at 5cm greater than the lowest snow depth value and vice versa for the 80% quantile and the highest snow depth value. These steps limited the ability of extreme outlier snow depths to influence the relationship between snow depth and reproduction. We used root-mean-squared error (RMSE) to compare the fit of ensemble averages built from all models in the ensemble versus only models in which either the pre- or post-threshold snow depth term (or the linear term if the linear model had the lowest AIC value) was a statistically significant predictor ($p<0.1$) of chickadees observed.

We explored how robust the relationship between snow depth and post-breeding counts of chickadees was to differences in the geographic regions and time frames used for filtering the eBird data. We used the same combinations of geographic regions and time frames as in our exploratory analysis in the first validation step, but here we used single models (with site ID as a random effect) to examine the relationship between snow depth and post-breeding chickadee counts in each dataset. Single models, instead of ensemble models, were a better method for this analysis because of the computational constraints of combining the ensemble approach with threshold models across many different datasets. This decision was justified by the fact that the single models produced relatively similar results as the ensemble approach. Our findings in this section provide confidence that the relationships between snow depth and reproduction are fairly robust to minor differences in the geographic range extents and time frames used to filter eBird data.

*Timing of mountain chickadee reproduction in the Sierra Nevada*

In the long-term study, low elevation first egg dates ranged from 3 May to 17 June (average 20 May), and high elevation first egg dates ranged from 19 May to 22 June (average 2 June). North of the long-term site, Dahlsten et al. (1992) report that mountain chickadees in Modoc county, CA initiated nests between 8 May and 13 June (approximate average 20 May; Figure 3 in Dhalsten et al.) at elevations lower than and equal to our low-elevation site. Far to the south of the long-term site, first-egg dates in the Sierra National Forest, CA ranged from approximately 7 May to 19 June at elevations that encompassed both our low and high sites (Coe et al. 2021). Finally, east of the Sierra range and southeast of Sierra National Forest, mountain chickadees in the White and Inyo Mountains initiated egg laying between 17 May and 23 May at elevations between our low and high sites (Hall and Morrison 2003). While limited, these separate observations suggest that timing of breeding is remarkably consistent among mountain chickadee populations throughout the Sierra Nevada, meaning the post-breeding periods we identified in the long-term study are likely capturing similar measures of post-breeding abundance throughout the Sierras.

**Supplemental figures**

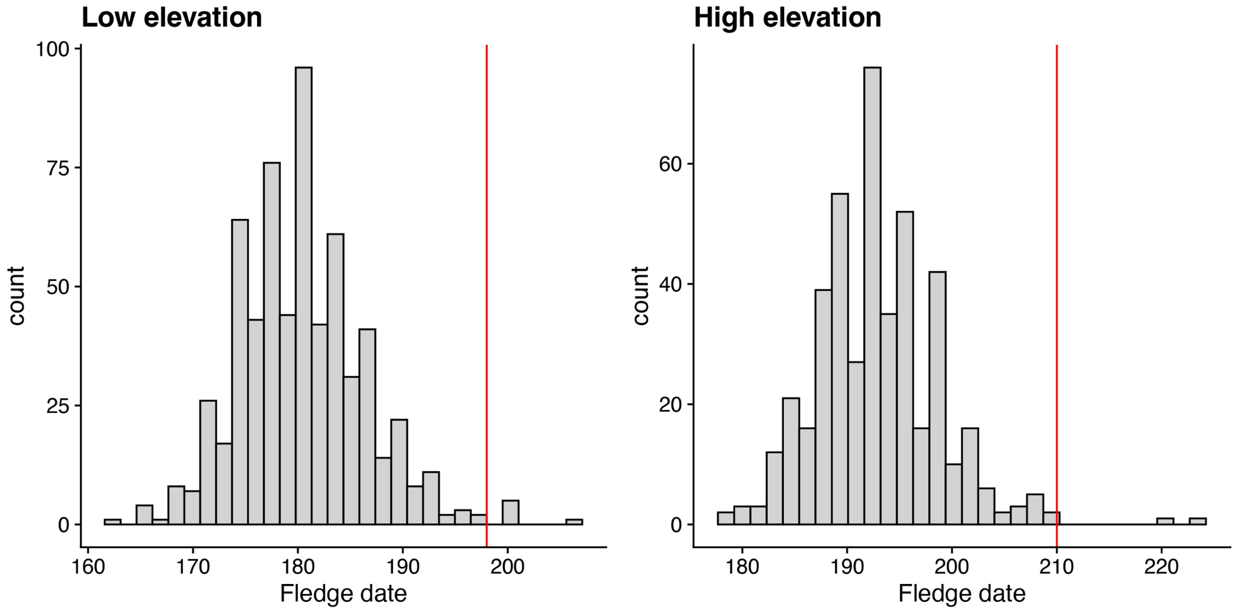


**Supplemental Figure S1**. Fledge dates (day of year) of mountain chickadee nests at the low and high elevation sites in the long-term study between 2014 and 2024. Red lines show the start date of the post-breeding period when 99% or more nests in the long-term study had fledged before these dates.

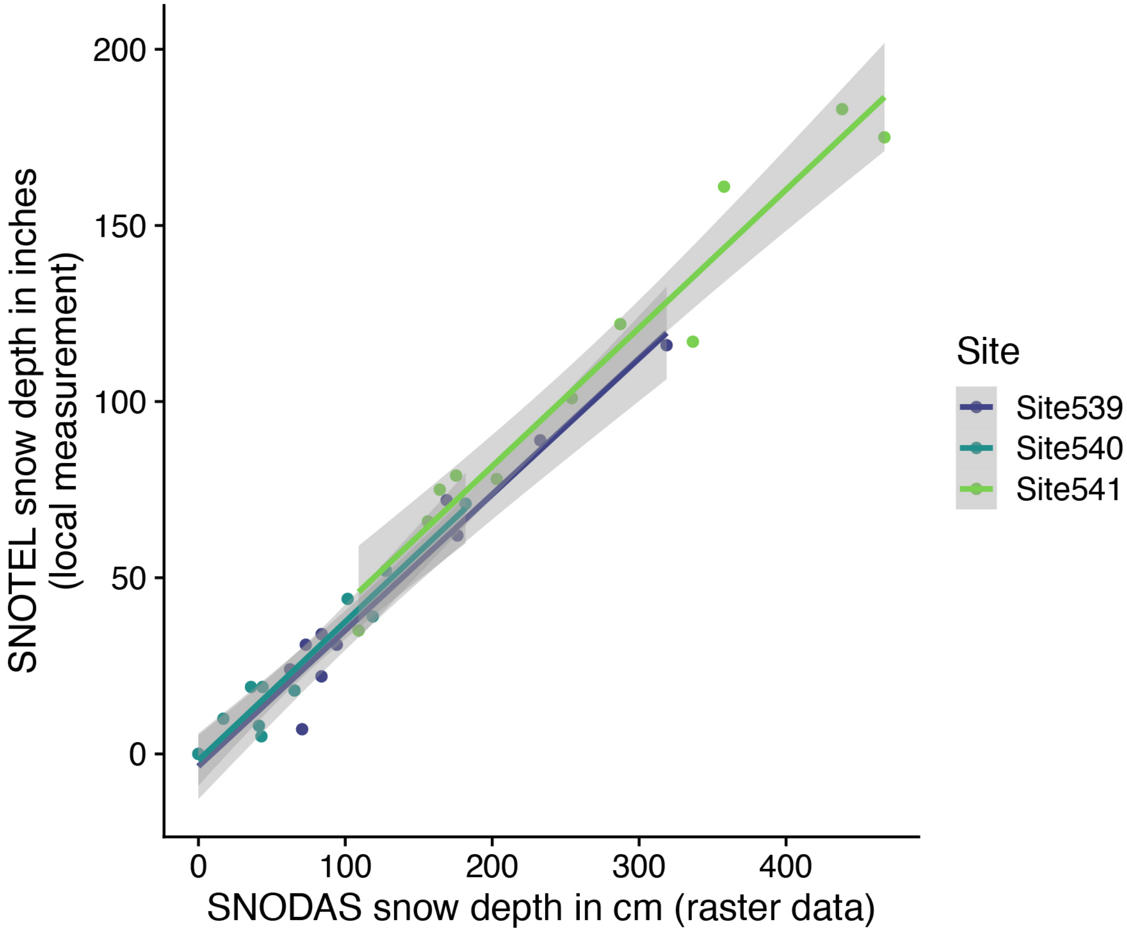


**Supplemental Figure S2**. Correlation between SNODAS (SNOw Data Assimilation System) snow depth data and SNOTEL (Snow Telemetry) snow depth data at the locations of three SNOTEL sites near the long-term study site. The two measurement methods are very highly correlated: Site 539 x SNODAS correlation adjusted $R^2$ = 0.95, Site 540 x SNODAS correlation adjusted $R^2$ = 0.92, Site 541 x SNODAS correlation adjusted $R^2$ = 0.95.

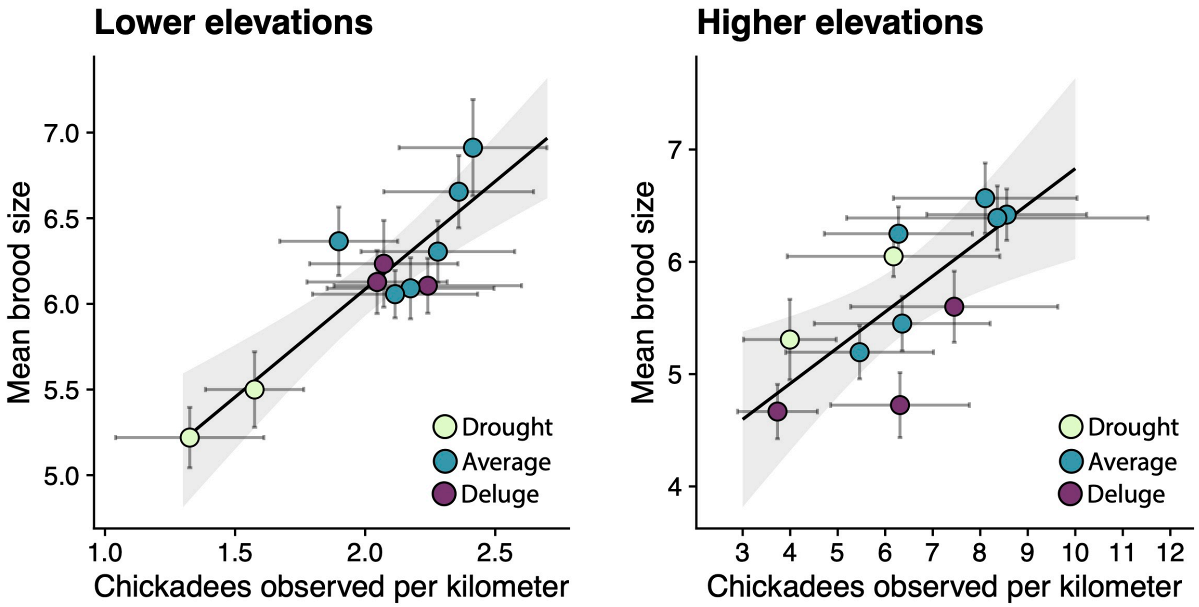


**Supplemental Figure S3**. Validation step 1: test of whether post-breeding eBird counts of chickadees can correlate with brood sizes from the long-term study site. A) Correlation of mean brood sizes at the low elevation site and post-breeding counts of chickadees in eBird data collected from July 17 to Sept 9 (end of the 36$^{th}$ week of the year) and from latitudes 39.2° to 39.6° N at lower elevations. B) Correlation of mean brood sizes at the high elevation site and post-breeding counts of chickadees in eBird data collected from July 29 to Aug 19 (end of the 33$^{rd}$ week of the year) and from latitudes 38.6° to 40.2° N at higher elevations. Error bars for chickadees per kilometer show 95% reference intervals, showing the approximate distribution of the predicted values for each year using the ensemble average approach.

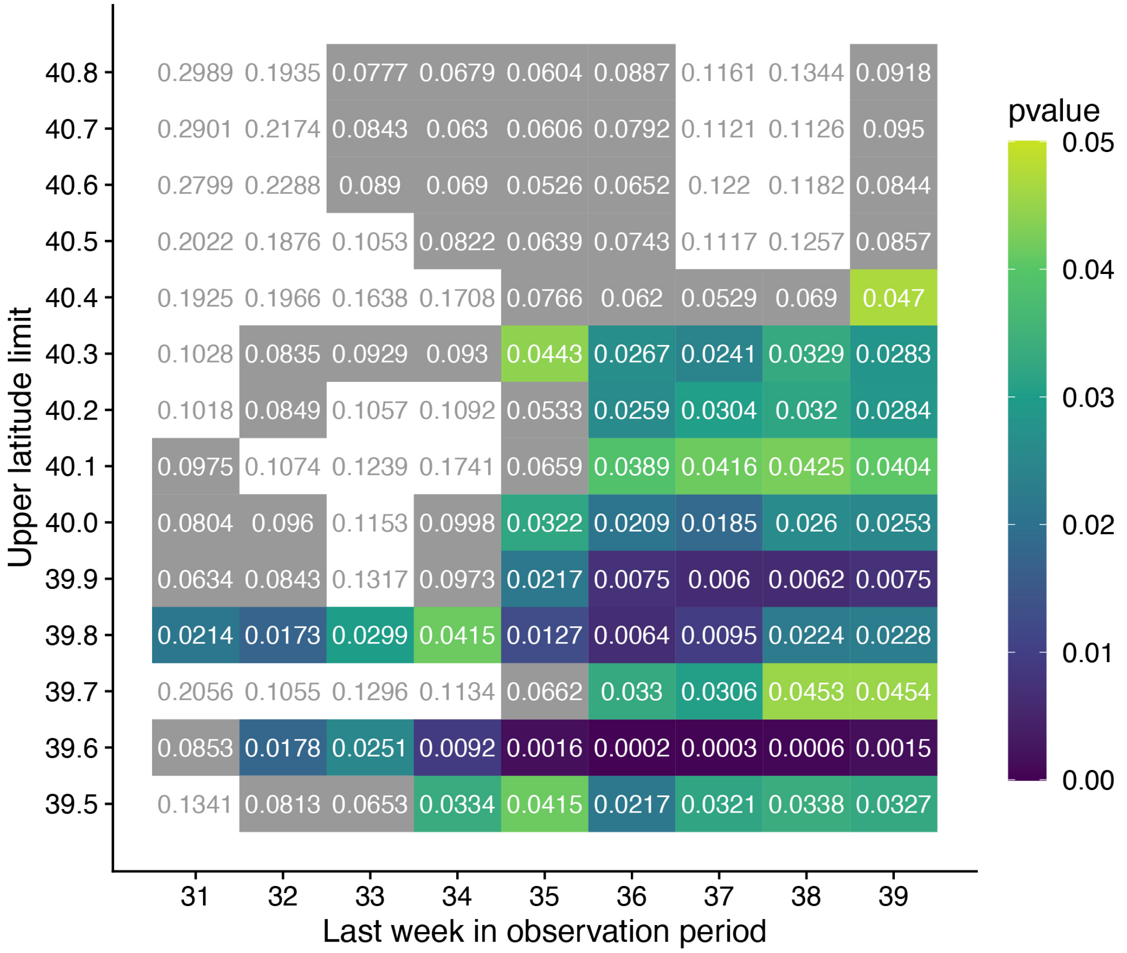


**Supplemental Figure S4**. Validation step 1: exploratory test of whether post-breeding eBird counts of chickadees collected between 1829m and 2134m (6000-7000ft) can correlate with brood sizes from the low-elevation site in the long-term data. Average yearly brood sizes in the long-term data were compared to ensemble average predictions of yearly post-breeding chickadee counts in eBird data from different time periods and geographic regions. X-axis: terminal week in the eBird data observation period. Each observation period started on day 198 on a 365-day calendar (the approximate last fledgling date at the low elevation site in the long-term study), so day 198 to the end of week 31 would represent a 21-day period. Y-axis: upper latitude limit of the geographic range for the eBird data. Latitude windows were centered on the latitude of the long-term field site (39.4° N) and extended in equal degrees to the north and south. A latitude window with an upper limit of 39.5° N would have a lower limit of 39.3° N. Each cell represents a different eBird dataset filtered for the corresponding x and y-axis parameters. Cell values show p-values from correlations between average yearly brood sizes at the low elevation site in the long-term data and yearly ensemble averages of chickadees observed per kilometer in the eBird data. Colored cells represent eBird datasets that correlated well with the long-term data ($p<0.05$), whereas cells with a gray background represent eBird datasets that correlated less-strongly with the long-term data ($p<0.10$, $p>0.05$). White cells did not correlate with the long-term data ($p>0.10$). The figure shows that certain time periods and latitude ranges in the eBird data are better matches to the long-term data than others, but the correlation to the long-term data is somewhat robust to different time periods and latitudes.

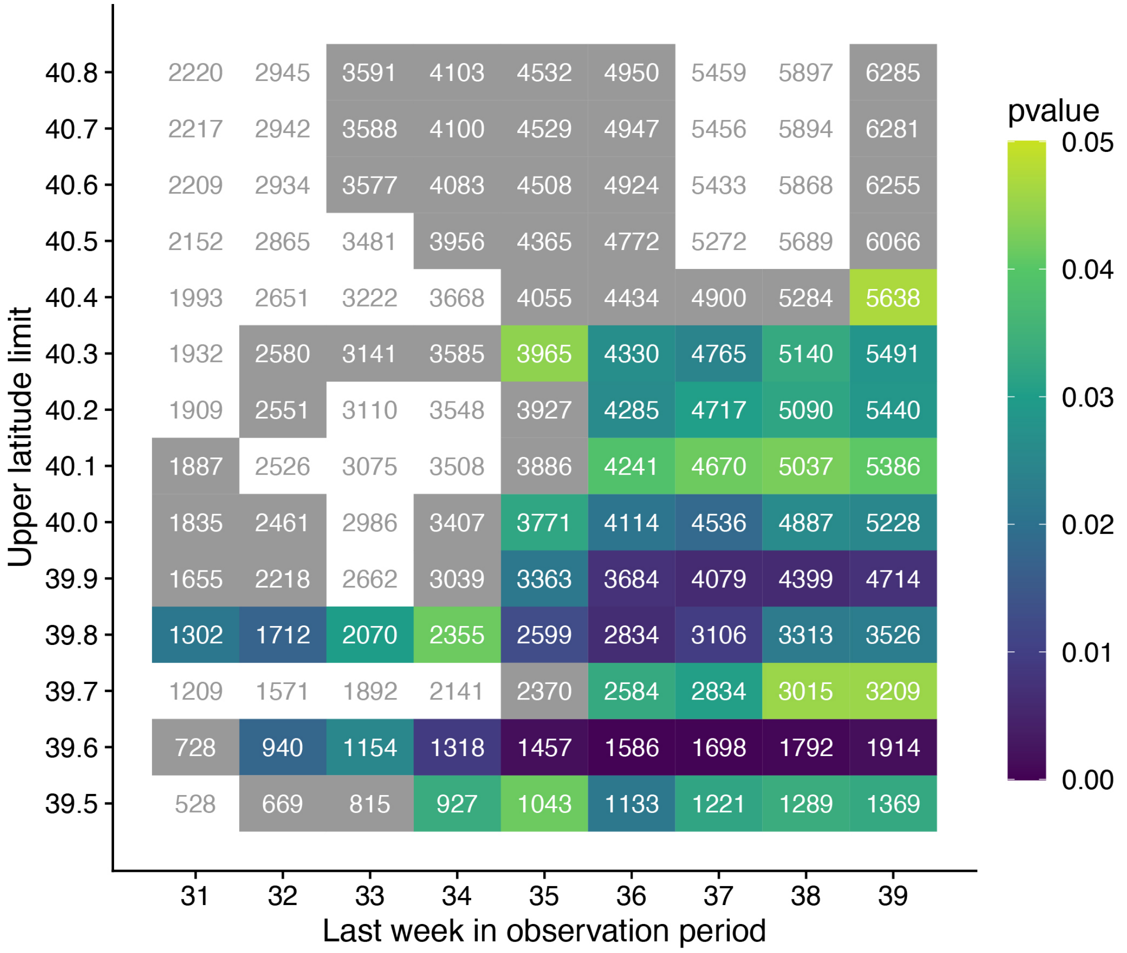


**Supplemental Figure S5**. Validation step 1: sample sizes from each eBird dataset associated with the above figure comparing the long-term data at the low elevation site to post-breeding chickadee counts in eBird data from different time periods and geographic regions. X-axis: terminal week in the eBird data observation period. Each observation period started on day 198 on a 365-day calendar (the approximate last fledgling date at the low elevation site in the long-term study), so day 198 to the end of week 31 would represent a 21-day period. Y-axis: upper latitude limit of the geographic range for the eBird data. Latitude windows were centered on the latitude of the long-term field site (39.4° N) and extended in equal degrees to the north and south. A latitude window with an upper limit of 39.5° N would have a lower limit of 39.3° N. Each cell represents a different eBird dataset filtered for the corresponding x and y-axis parameters. Cell values show sample sizes from each eBird dataset. Colored cells represent eBird datasets that correlated well with the long-term data ($p<0.05$), whereas cells with a gray background represent eBird datasets that correlated less-strongly with the long-term data ($p<0.10$, $p>0.05$). White cells did not correlate with the long-term data ($p>0.10$).

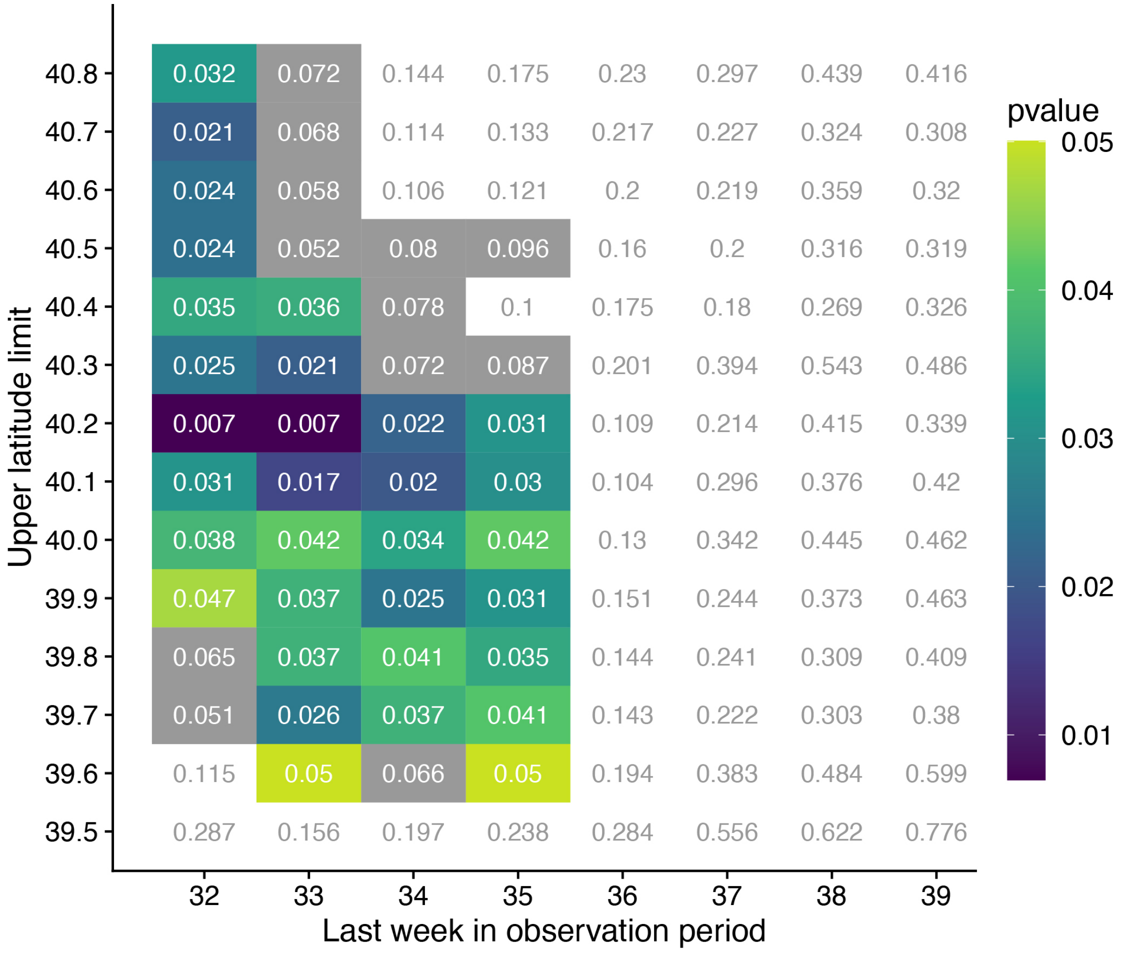


**Supplemental Figure S6**. Validation step 1: exploratory test of whether post-breeding eBird counts of chickadees collected from elevations above 2377m (7800ft) can correlate with brood sizes from the high-elevation site in the long-term data. Average yearly brood sizes in the long-term data were compared to ensemble average predictions of yearly post-breeding chickadee counts in eBird data from different time periods and geographic regions. X-axis: terminal week in the eBird data observation period. Each observation period started on day 210 on a 365-day calendar (the approximate last fledgling date at the high elevation site in the long-term study), so day 210 to the end of week 32 would represent a 15-day period. Y-axis: upper latitude limit of the geographic range for the eBird data. Latitude windows were centered on the latitude of the long-term field site (39.4° N) and extended in equal degrees to the north and south. A latitude window with an upper limit of 39.5° N would have a lower limit of 39.3° N. Each cell represents a different eBird dataset filtered for the corresponding x and y-axis parameters. Cell values show p-values from correlations between average yearly brood sizes at the high elevation site in the long-term data and yearly ensemble averages of chickadees observed per kilometer in eBird data. Colored cells represent eBird datasets that correlated well with the long-term data ($p<0.05$), whereas cells with a gray background represent eBird datasets that correlated less-strongly with the long-term data ($p<0.10$, $p>0.05$). White cells did not correlate with the long-term data ($p>0.10$). The figure shows that certain time periods and latitude ranges in the eBird data are better matches to the long-term data than others, but the correlation to the long-term data is somewhat robust to different time periods and latitudes.

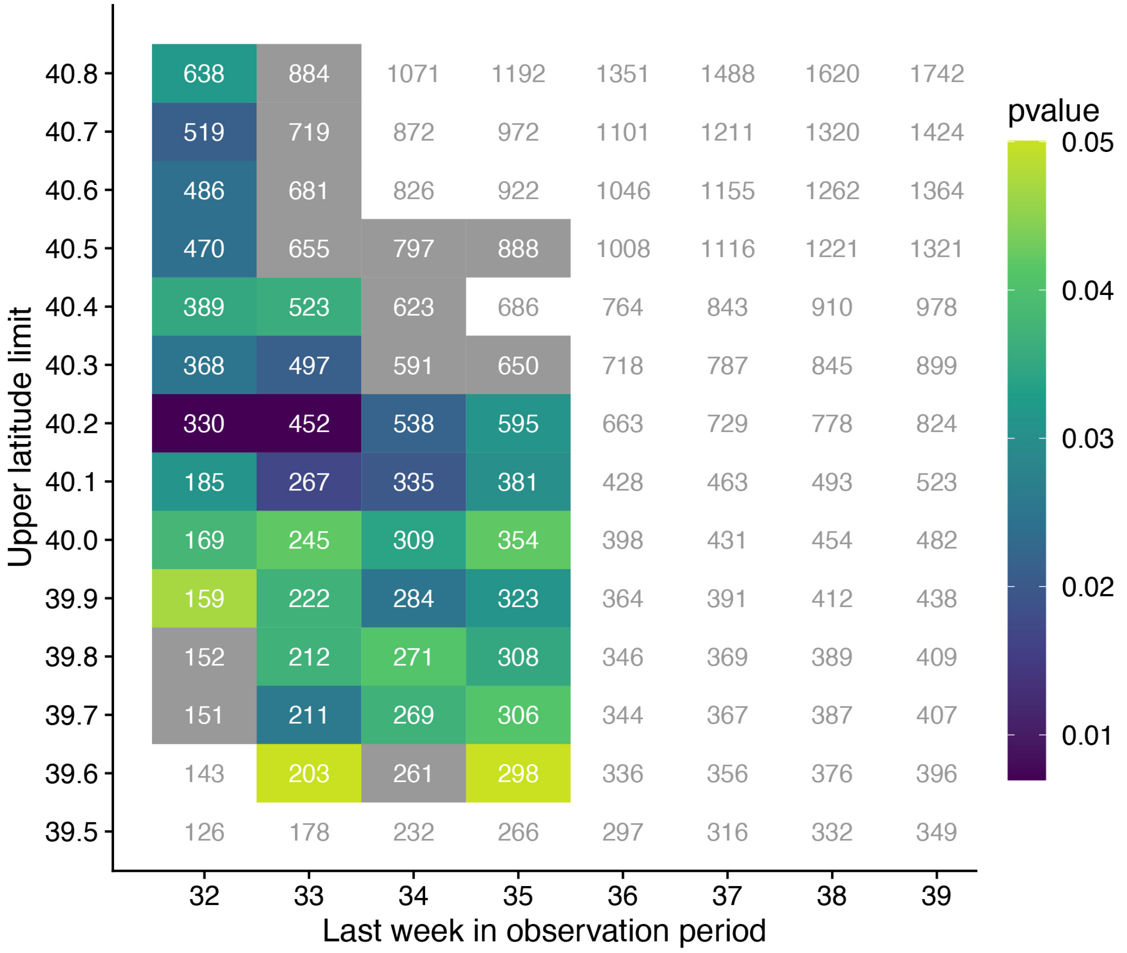


**Supplemental Figure S7**. Validation step 1: sample sizes from each eBird dataset associated with the above figure comparing the long-term data at the high-elevation site to post-breeding chickadee counts in eBird data from different time periods and geographic regions. X-axis: terminal week in the eBird data observation period. Each observation period started on day 210 on a 365-day calendar (the approximate last fledgling date at the low elevation site in the long-term study), so day 210 to the end of week 32 would represent a 15-day period. Y-axis: upper latitude limit of the geographic range for the eBird data. Latitude windows were centered on the latitude of the long-term field site (39.4° N) and extended in equal degrees to the north and south. A latitude window with an upper limit of 39.5° N would have a lower limit of 39.3° N. Each cell represents a different eBird dataset filtered for the corresponding x and y-axis parameters. Cell values show sample sizes from each eBird dataset. Colored cells represent eBird datasets that correlated well with the long-term data ($p<0.05$), whereas cells with a gray background represent eBird datasets that correlated less-strongly with the long-term data ($p<0.10$, $p>0.05$). White cells did not correlate with the long-term data ($p>0.10$).

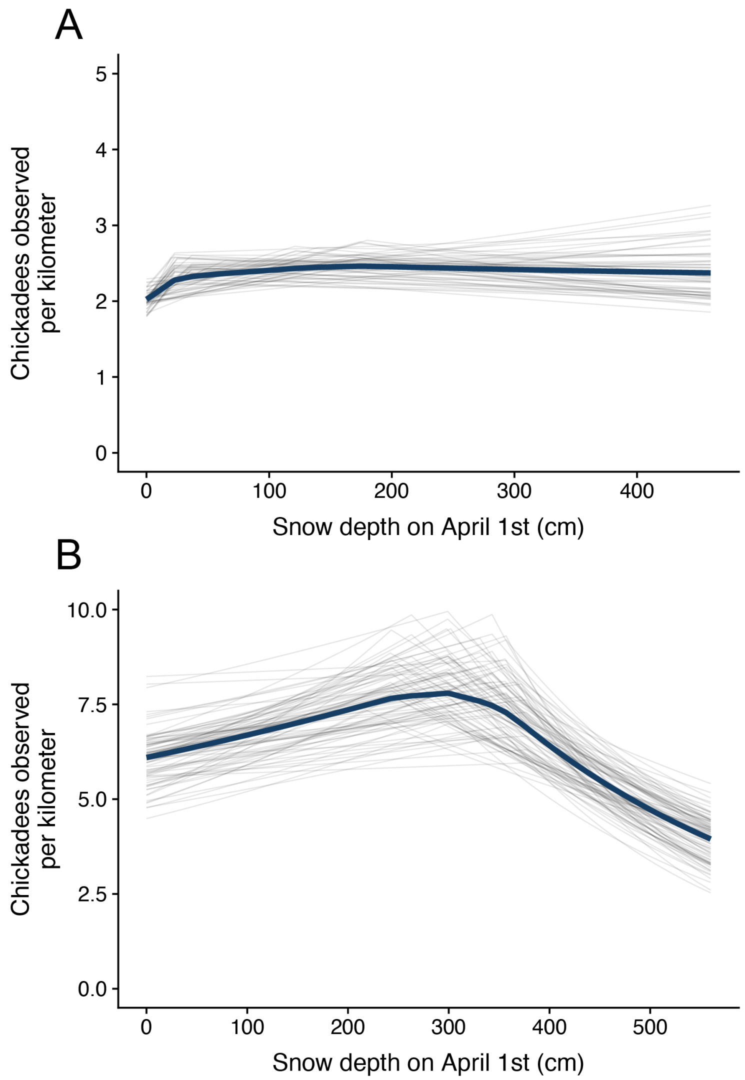


**Supplemental Figure S8**. Validation step 2: testing whether the eBird datasets that best correlated with the long-term data show similar relationships between snow depth and reproduction as seen in the long-term data. Model predictions calculated using the ensemble average method are shown (dark lines). Light gray lines show the individual model predictions. A) Ensemble average prediction for the relationship between snow depth and mountain chickadee counts in eBird data near the long-term study site from elevations between 1829m and 2134m (6000-7000ft), corresponding to the low-elevation site in the long-term study. The eBird dataset included checklists collected between latitudes 39.2° N and 39.6° N and from day 198 and through week 36. B) Ensemble average prediction for the relationship between snow depth and mountain chickadee counts in eBird data near the long-term study site from elevations equal to or above 2377m (7800ft), corresponding to the high-elevation site in the long-term study. The eBird dataset included checklists collected between latitudes 38.6° N and 40.2° N and from day 210 through week 33.

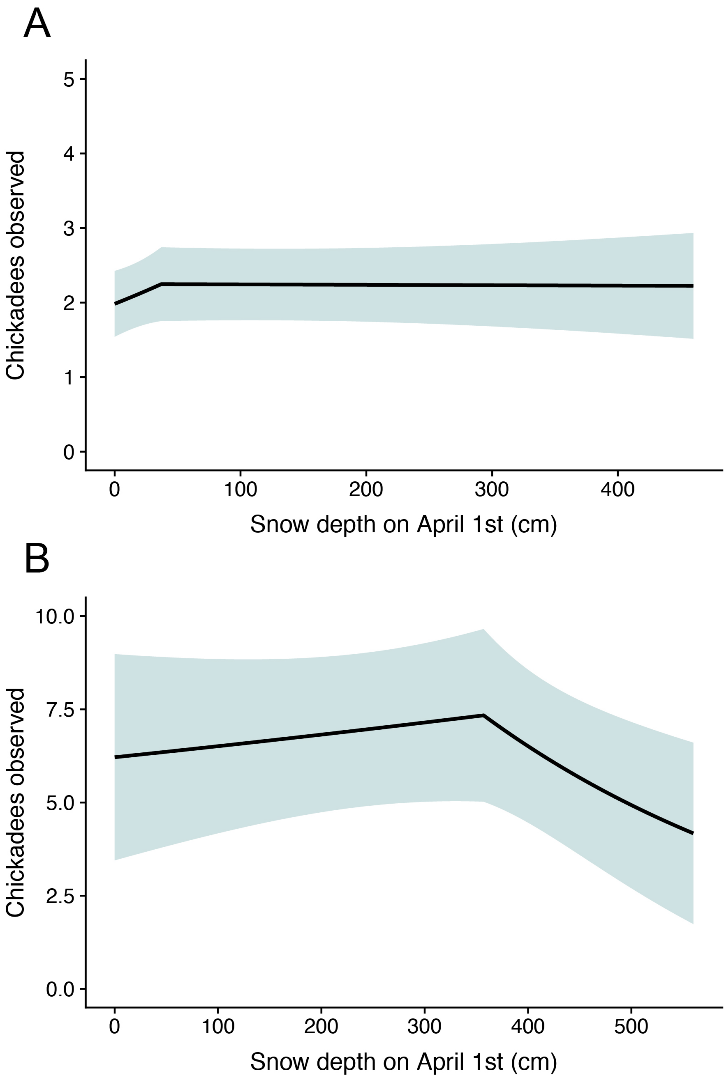


**Supplemental Figure S9**. Validation step 2: testing whether the eBird datasets that best correlated with the long-term data show similar relationships between snow depth and reproduction as seen in the long-term data. Model predictions for single models, which included checklist location as a random effect, are shown. A) Single model prediction for the relationship between snow depth and mountain chickadee counts in eBird data near the long-term study site from elevations between 1829m and 2134m (6000-7000ft), corresponding to the low-elevation site in the long-term study. The eBird dataset included checklists collected between latitudes 39.2° N and 39.6° N and from day 198 and through week 36. B) Single model prediction for the relationship between snow depth and mountain chickadee counts in eBird data near the long-term study site from elevations equal to or above 2377m (7800ft), corresponding to the high-elevation site in the long-term study. The eBird dataset included checklists collected between latitudes 38.6° N and 40.2° N and from day 210 through week 33.

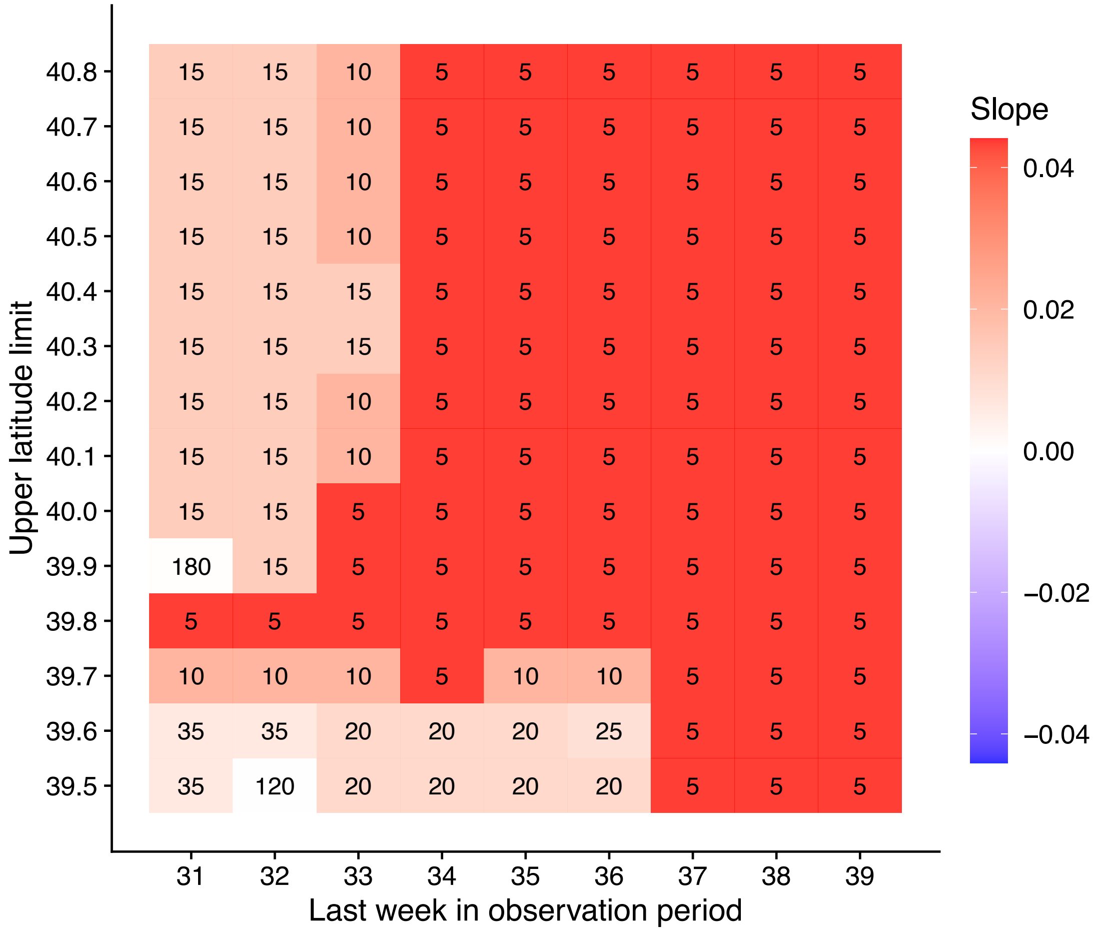


**Supplemental figure S10**. Validation step 2: testing the robustness of the relationship between snow depth and reproduction in eBird data collected between 1829m and 2134m (6000-7000ft), corresponding to the low-elevation site in the long-term study. X-axis: terminal week in the eBird data observation period. Each observation period started on day 198 on a 365-day calendar (the approximate last fledgling date at the low elevation site in the long-term study), so day 198 to the end of week 31 would represent a 21-day period. Y-axis: upper latitude limit of the geographic range for the eBird data. Latitude windows were centered on the latitude of the long-term field site (39.4° N) and extended in equal degrees to the north and south. A latitude window with an upper limit of 39.5° N would have a lower limit of 39.3° N. Each cell represents a different eBird dataset filtered for the corresponding x and y-axis parameters. Cell numbers come from models investigating the relationship between snow depth and chickadees observed per kilometer in each eBird dataset and represent the threshold value for the snow depth term (snow depth value at which the slope changed) that was best supported by AIC. Cell colors correspond to the pre-threshold slope in this figure, with red being positive and blue being negative slopes. The figure shows that the relationship between snow depth and chickadees observed per kilometer in eBird data at elevations between 1829m and 2134m (6000-7000ft) is similar across geographic regions and observation periods. Nearly all datasets explored exhibited a positive slope before the threshold value, matching the prediction of the eBird dataset that best correlated with the long-term data (Supplemental figure S7A, S8A).

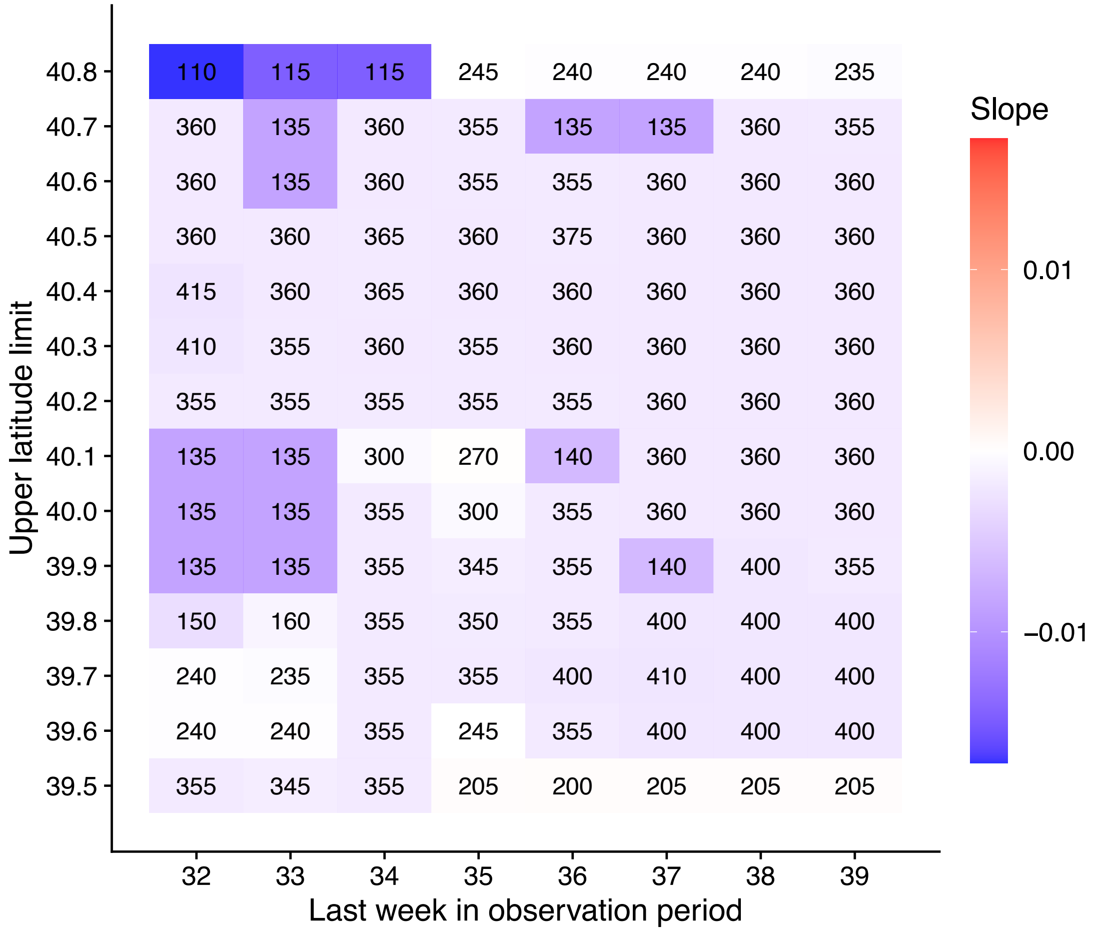


**Supplemental figure S11**. Validation step 2: testing the robustness of the relationship between snow depth and reproduction in eBird data collected above 2377m (7800ft), corresponding to the high-elevation site in the long-term study. X-axis: terminal week in the eBird data observation period. Each observation period started on day 210 on a 365-day calendar (the approximate last fledgling date at the high elevation site in the long-term study), so day 210 to the end of week 32 would represent a 15-day period. Y-axis: upper latitude limit of the geographic range for the eBird data. Latitude windows were centered on the latitude of the long-term field site (39.4° N) and extended in equal degrees to the north and south. A latitude window with an upper limit of 39.5° N would have a lower limit of 39.3° N. Each cell represents a different eBird dataset filtered for the corresponding x and y-axis parameters. Cell numbers come from models investigating the relationship between snow depth and chickadees observed per kilometer in each eBird dataset and represent the threshold value for the snow depth term (snow depth value at which the slope changed) that was best supported by AIC. Cell colors correspond to the post-threshold slope in this figure, with red being positive and blue being negative slopes. The figure shows that the relationship between snow depth and chickadees observed per kilometer in eBird data at elevations above 2377m (7800ft) is similar across geographic regions and observation periods. Nearly all datasets explored exhibited a negative slope after the threshold value, matching the prediction of the eBird dataset that best correlated with the long-term data (Supplemental figure S7B, S8B).

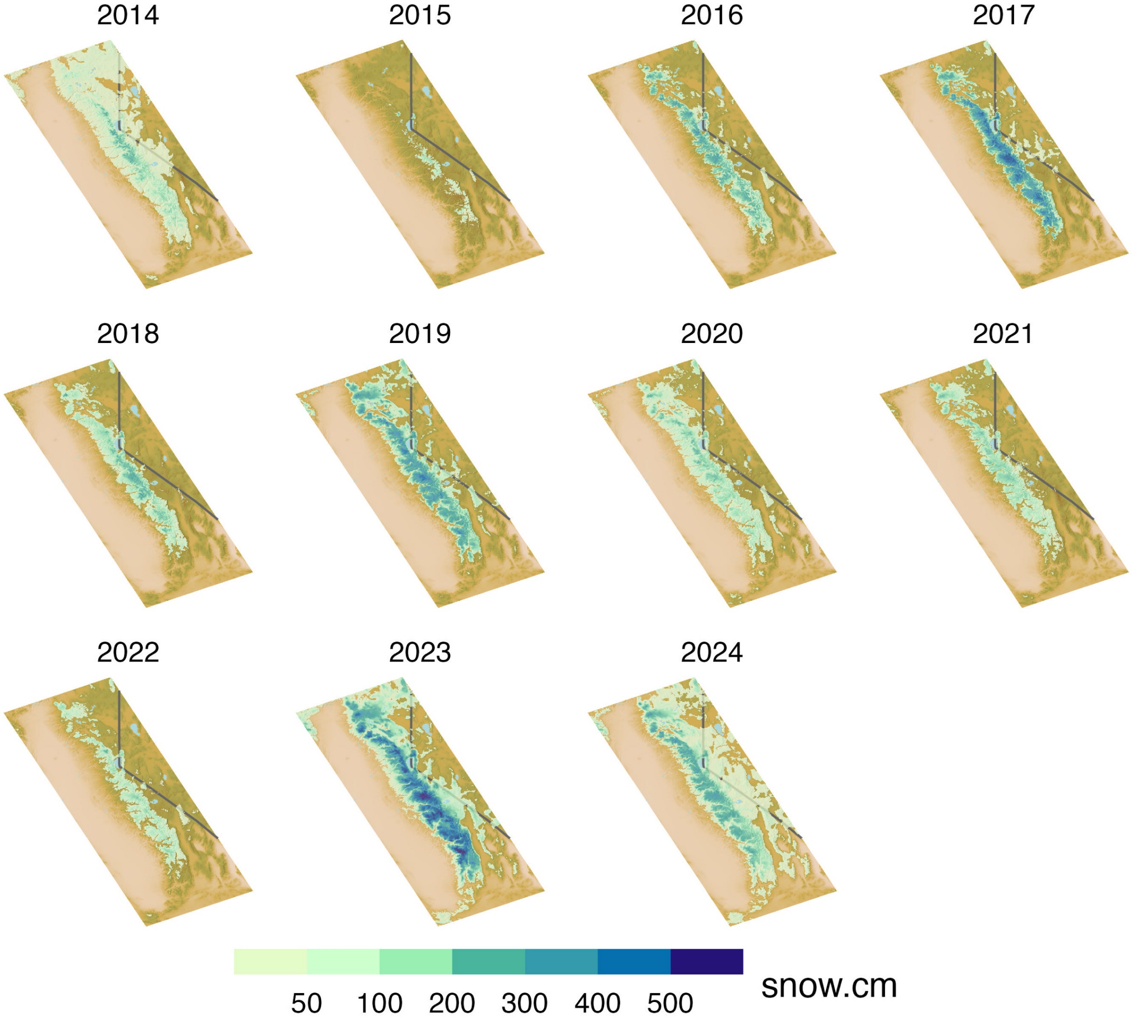


**Supplemental figure S12**. Maps of Sierra Nevada snow depth on April 1st by year.

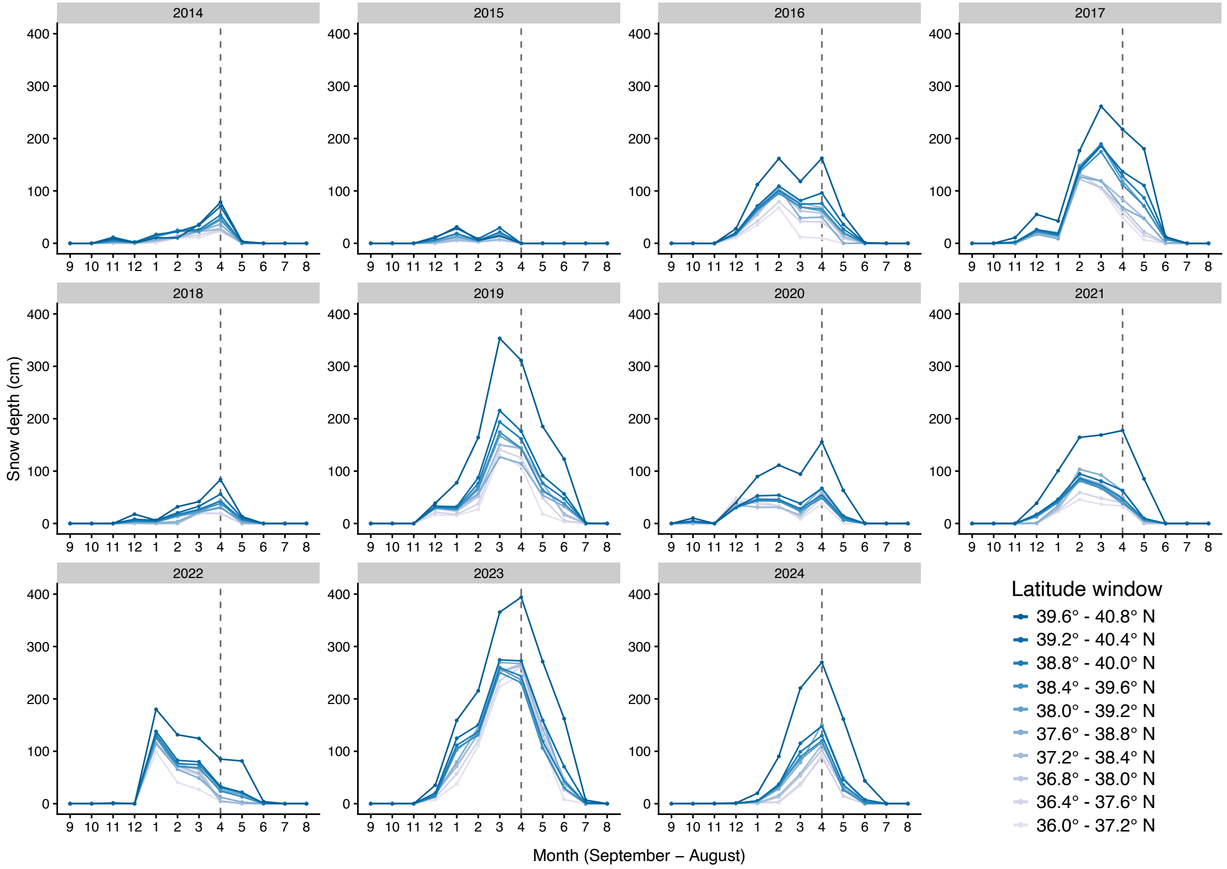


**Supplemental figure S13**. Average monthly snow depth at eBird checklist locations by latitude window and year from 2014 – 2024 for eBird checklist locations collected at elevations between 1829m and 2134m (6000-7000ft), corresponding to the low-elevation site in the long-term study.

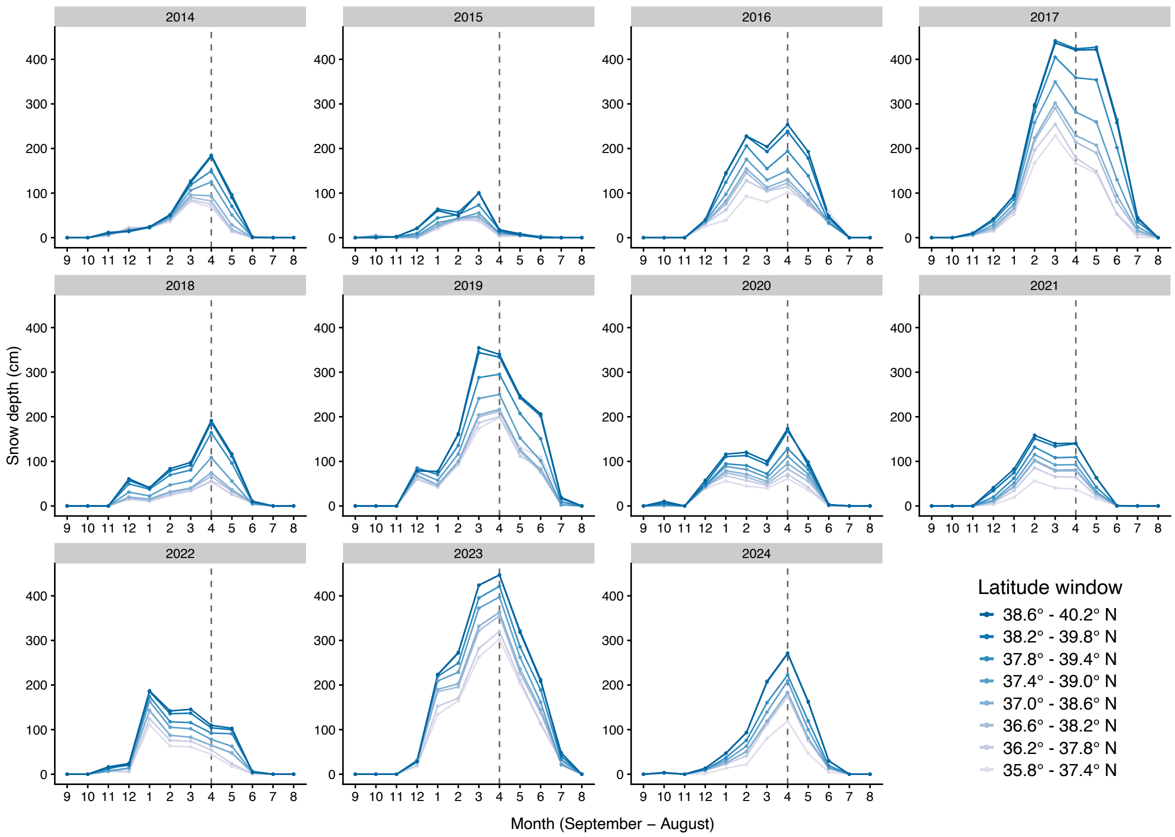


**Supplemental figure S14**. Average monthly snow depth at eBird checklist locations by latitude window and year from 2014 – 2024 for eBird checklist locations collected at elevations above 2377m (7800ft), corresponding to the high-elevation site in the long-term study.
